# Targeted substrate degradation by Kelch controls the actin cytoskeleton during ring canal expansion

**DOI:** 10.1101/315077

**Authors:** Andrew M. Hudson, Katelynn M. Mannix, Julianne A. Gerdes, Molly C. Kottemann, Lynn Cooley

## Abstract

During Drosophila oogenesis, specialized actin-based structures called ring canals form and expand to accommodate growth of the oocyte. Previous work demonstrated that Kelch and Cullin 3 function together in a Cullin 3-RING ubiquitin ligase complex (CRL3^Kelch^) to organize the ring canal cytoskeleton, presumably by targeting a substrate for proteolysis. Here, we use tandem affinity purification followed by mass spectrometry to identify HtsRC as the CRL3^Kelch^ ring canal substrate. CRISPR-mediated mutagenesis of HtsRC revealed its requirement in the recruitment of the ring canal F-actin cytoskeleton. We present genetic evidence consistent with HtsRC being the CRL3^Kelch^ substrate, as well as biochemical evidence indicating that HtsRC is ubiquitylated and degraded by the proteasome. Finally, we identify a short sequence motif in HtsRC that is necessary for Kelch binding. These findings uncover an unusual mechanism during development wherein a specialized cytoskeletal structure is regulated and remodeled by the ubiquitin-proteasome system.

## Introduction

Gamete formation in animals results in the production of the highly specialized cells responsible for species propagation, the sperm and egg (Matova and Cooley, 2001; Roosen-Runge, 1969). The cell biology involved in gametogenesis is in many instances unusual: cells often develop in interconnected cysts, mature sperm have tails up to millimeters in length, oocytes can grow to macroscopic sizes, and the apoptotic machinery is used extensively both to promote waves of canonical apoptosis and also to coordinate non-apoptotic development and differentiation (Arama et al., 2003; Peterson et al., 2015). Many questions remain about how basic cell biological mechanisms are coordinated to achieve the remarkable form and function of animal gametes.

Drosophila oogenesis provides a powerful model for investigating the specialized features of gamete development (Bastock and St Johnston, 2008). Female fruit flies produce up to 60 eggs per day, each approximately 0.5 mm in length with a volume more than 1000-fold greater than that of a typical somatic cell. This level of fecundity is achieved through mass production: the two Drosophila ovaries are made up of approximately 16 ovarioles, each of which contains a linear array of progressively developing eggs. Oogenesis initiates at the anterior end of an ovariole, called the germarium, where a daughter of a germline stem cell undergoes four rounds of mitosis to generate a 16-cell cyst. One of these cells becomes the oocyte, while the remaining 15 differentiate as nurse cells. Cytokinesis does not complete during the mitotic divisions, leaving all 16 cells connected by the arrested cleavage furrows. The arrested cleavage furrows are subsequently transformed into stable intercellular bridges called ring canals through the recruitment of additional proteins (Haglund et al., 2011). Over the course of several days, the oocyte grows in volume by five orders of magnitude without a substantial transcriptional contribution from the oocyte nucleus (Cummings et al., 1971). Instead, the nurse cells become highly polyploid and synthesize mRNA, protein, and organelles that support oocyte growth. These components move to the oocyte through the ring canals, which must also grow to accommodate the flux of materials. Ring canals recruit a robust F-actin cytoskeleton and expand during oogenesis, reaching diameters greater than 10 *μ*m.

An important mechanism for regulating developmental events during gametogenesis is the ubiquitin-proteasome system (UPS). Covalent attachment of the small protein ubiquitin to a target protein can alter the fate of the target protein in a number of ways (Komander and Rape, 2012). Monoubiquitylation, the attachment of a single ubiquitin, can serve as a signal for endocytic sorting. When a chain of ubiquitins linked in series though ubiquitin Lysine 48 (K48-linked chains) are assembled, the ubiquitylated protein is targeted for destruction by the proteasome. Protein ubiquitylation is carried out by three enzymes acting in sequence, termed E1, E2, and E3. Substrate specificity is conferred by the E3 enzyme, and several classes of E3 enzymes have been characterized. Genes encoding E3 enzymes have undergone significant expansion in higher eukaryotes such that 3% of human genes (>600 out of 20k) encode substrate-specific E3 enzymes (Li et al., 2008).

BTB-BACK-Kelch (BBK) proteins function as substrate adaptors for a class of ubiquitin E3 ligases called Cullin 3-RING ubiquitin ligases (CRL3s) (Figure 1A) (Xu et al., 2003). The elongated Cullin 3 (Cul3) protein binds a BTB-domain protein at its N-terminus and a RING domain protein at its C-terminus. The RING domain recruits an E2 ubiquitin conjugating enzyme, while the BTB-domain protein recruits specific substrates for ubiquitylation through their C-terminal Kelch-repeat domains (KREP), which adopt a *β*-propeller structure (Adams et al., 2000; Li et al., 2004). The BBK gene family has undergone significant expansion during animal evolution; the Drosophila genome encodes ~10 BBK proteins, while nearly 50 distinct BBK genes are present in humans (Dhanoa et al., 2013; Prag and Adams, 2003). Mutations in human BBK genes are associated with a number of diseases including cancer, neurodegenerative disease, infertility, and congenital skeletal myopathies, underscoring their clinical importance (Bomont et al., 2000; Gupta and Beggs, 2014; Padmanabhan et al., 2006; Yatsenko et al., 2006). A common thread suggested by these disease associations is BBK-mediated regulation of metazoan-specific cytoskeletal structures. Studying the targets of BBK/CRL3 ubiquitylation therefore requires access to experimental systems amenable to analysis of cytoskeletal regulation.

We previously showed that Drosophila Kelch functions with Cullin 3 to coordinate the growth of the ovarian ring canal cytoskeleton during oogenesis (Hudson and Cooley, 2010; Hudson et al., 2015). We provided genetic evidence that Kelch, Cullin 3, and the proteasome function in a common pathway targeting one or more substrates for ubiquitylation and degradation. Here, we present evidence showing that HtsRC, a critical regulator of F-actin recruitment to ovarian ring canals, is the substrate of CRL3^Kelch^ during oogenesis. We show that the Kelch KREP domain and HtsRC can be detected in a complex, and we define a linear sequence motif in HtsRC responsible for targeting by CRL3^Kelch^. We also describe a series of genetic experiments consistent with HtsRC being the ring canal substrate of CRL3^Kelch^, and finally demonstrate that HtsRC can be ubiquitylated and degraded by the proteasome in cultured cells.

## Results

### Tandem affinity purification and mass spectrometry reveals HtsRC as a potential CRL3^Kelch^ substrate

Egg chambers from Drosophila females mutant for kelch have ring canals with a highly disorganized F-actin cytoskeleton that impedes the flow of cytoplasm to growing oocytes (Xue and Cooley, 1993). We have reported that Kelch functions as the substrate-targeting component of a Cullin-RING E3 ubiquitin ligase (CRL3^Kelch^) and that its CRL activity is essential for ring canal cytoskeletal organization (Hudson and Cooley, 2010; Hudson et al., 2015). Given these results, we looked for candidate substrates of CRL3^Kelch^ using an affinity purification / mass spectrometry (AP-MS) strategy. We previously showed that overexpression of the KREP domain results in a *kelch*-like dominant-negative phenotype (Figure 1C), marked by the accumulation of the KREP domain at the ring canal (Hudson and Cooley, 2010). This phenotype suggested that the KREP domain might be bound to its substrate in a stable complex since the KREP domain alone cannot associate with Cul3 and consequently should not promote substrate ubiquitylation. We therefore used affinity purification of the KREP domain to identify potential substrates by mass spectrometry.

To identify KREP-associated proteins, we expressed TAP-tagged mCherry::KREP (Figure 1B) and a TAP-tagged mCherry control construct in ovarian germ cells. However, initial experiments were hindered by the poor solubility of ring canal components lysed under conditions that would preserve protein-protein interactions. We therefore performed purifications in a mutant background to liberate ring canal components from the insoluble cytoskeleton. *cheerio* (*cher*) encodes Drosophila Filamin, the earliest known component recruited to stable germline ring canals (Sokol and Cooley, 1999). In *cher* mutants, ring canals lack the robust F-actin cytoskeleton found in wild type (Figures 1C and 1D), and the ring canal proteins HtsRC and Kelch fail to properly localize despite being present in ovarian lysates (Robinson et al., 1997). We performed parallel purifications of mCherry or mCherry::KREP from *cher* mutant females (Figure 1E) and identified co-purifying proteins by mass spectroscopy. From nearly 900 proteins identified in the KREP purification, we focused on 141 candidate substrates based on specificity of interaction and relative abundance (approximated by total spectral counts; see Table S1).

Sequence analysis of the candidate substrates revealed that they were enriched for proteins containing a short linear sequence motif consisting of several acidic residues followed by a consensus PEAEQ sequence (15 out of 141; P = 8 × 10^−8^, binomial test) (Figure 1F). A similar motif is also present in three human WNK kinases that are targeted by KLHL3, a human ortholog of Kelch (Boyden et al., 2012; Shibata et al., 2013). A peptide containing the PEADQ sequence derived from WNK4 binds KLHL3 protein in vitro, and the structure of a complex between KLHL3 and an acidic EPEEPEADQ peptide revealed that the peptide bound the ‘top’ surface of the KLHL3 KREP domain (Schumacher et al., 2014). These results suggest that the PEAEQ motif is a binding site for Kelch and closely related BBK proteins. In further support of this idea, we identified four PEAEQ-containing proteins in a yeast two-hybrid screen for KREP domain interactors, and all cDNAs isolated contained the regions encoding the PEAEQ motif (Figure S1).

**Figure 1.**
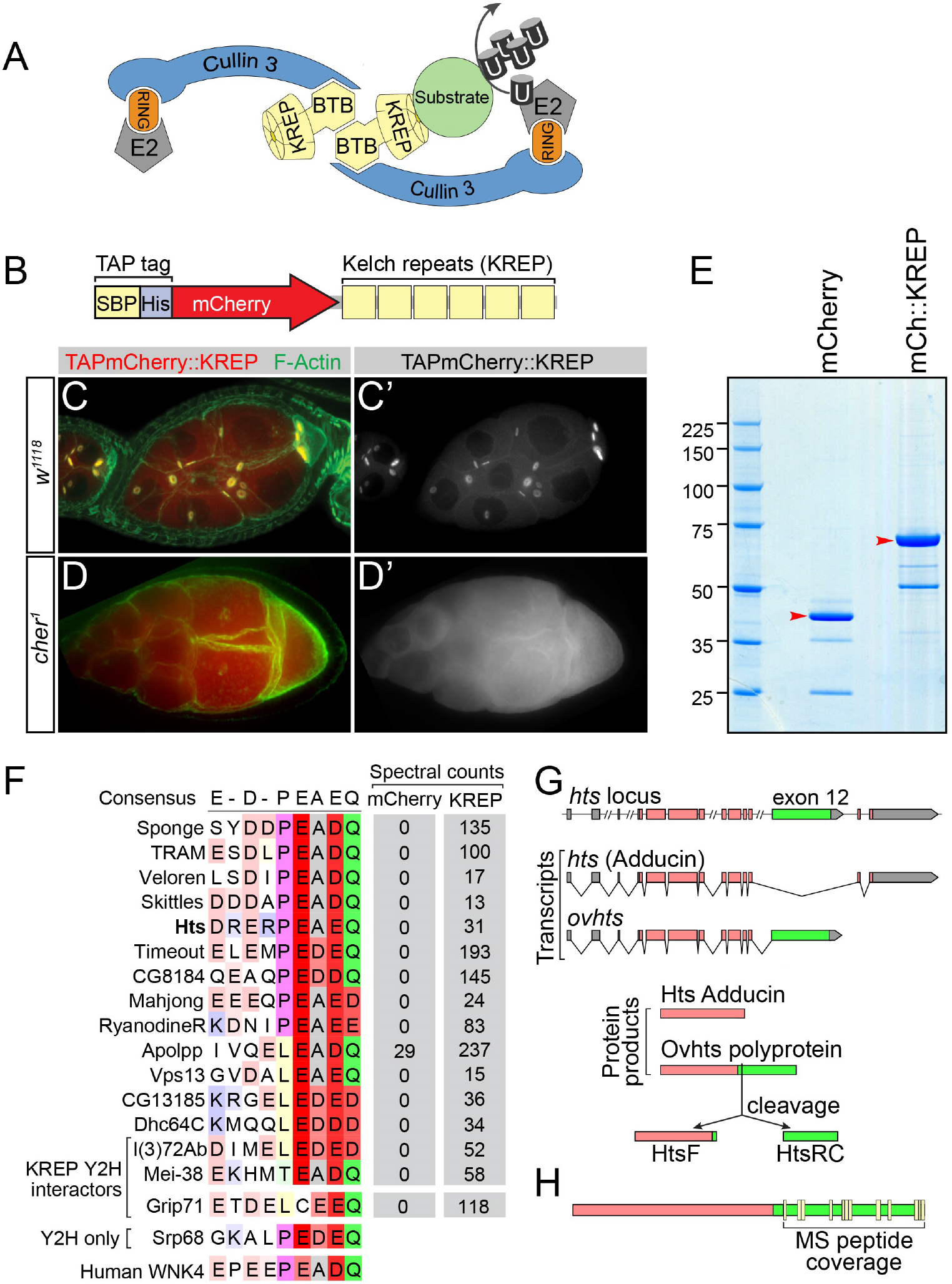
Tandem affinity purification and mass spectrometry reveals HtsRC as a potential CRL3^Kelch^ substrate. (A) Cartoon of a CRL3^Kelch^ dimer, showing ubiquitylation of a substrate (green circle). (B) Diagram of TAP-tagged mCherry fused to the Kelch KREP domain. (C-C″) High-level expression of the mCherry::KREP fusion using the *matGal4* driver results in ring canal localization and a dominant *kelch*-like phenotype. (D-D″) In *cher*^1^ mutant ovaries, the mCherry::KREP fusion exhibits a homogeneous cytosolic distribution and does not associate with ring canal cytoskeleton. (E) Proteins bound to TAP-tagged mCherry or TAPmCherry::KREP were purified from *cher*^1^ mutant ovaries, separated on a 4-12% SDS-PAGE gel, stained with colloidal coomassie, and subjected to LC-MS/MS. Purified TAPmCherry and TAPmCherry::KREP bands are indicated by red arrowheads. (F) Sequence analysis of proteins that bound specifically to TAPmCherry::KREP fusion protein identified a common motif consisting of several acidic residues followed by a PEAEQ consensus sequence defined by the expression [PLT]-E-[AD]-[DE]-[QD]. Sequences were manually aligned in JalView (Waterhouse et al., 2009). (G) Alternative splicing of *hts* transcripts produces several mRNAs, including mRNAs encoding Adducin. The ovary-specific *ovhts* mRNA contains exon 12 (green) and produces a polyprotein that is cleaved to produce truncated Adducin (HtsF) that localizes to fusomes and the novel HtsRC protein that localizes to ring canals. (H) Peptides identified from proteins bound to mCherry::KREP mapped entirely to HtsRC (yellow boxes). See also Figures S1, S2, and Table S1.

Several PEAEQ proteins identified by MS have known functions in F-actin assembly and organization, including Hts (Figure 1F). The *hts* locus encodes multiple proteins, including conserved and ubiquitously expressed isoforms of the cytoskeletal protein Adducin (Figure 1G) (Yue and Spradling, 1992). In the ovary, a germline-specific transcript designated *ovhts* encodes a polyprotein that is cleaved to produce two distinct products: an Adducin isoform that localizes to the fusome within mitotic cells (referred to as HtsF), and the novel Hts ring canal (HtsRC) polypeptide that functions at germline ring canals (Figure 1G) (Petrella et al., 2007). All of the Hts peptides identified by MS mapped to the HtsRC-encoding part of *ovHts* (Figure 1H), suggesting that the ring canal protein, and not an Adducin isoform, was associated with the KREP domain. Sequence analysis of other Drosophila species revealed that the HtsRC PEAEQ sequence is invariant, while adjacent sequences show less conservation, suggesting a functional role for this sequence (Figure S2). In addition, the association between the KREP domain and HtsRC in lysates prepared from *cher* mutants suggests that the interaction may be direct, and not the consequence of the two proteins being present together in a complex of ring canal proteins. We therefore carried out genetic and molecular experiments designed to determine whether HtsRC is a substrate of CRL3^Kelch^.

### Genetic analysis indicates *hts* is epistatic to *kelch*

Functional analyses of HtsRC have been limited due to the multiple requirements for hts during oogenesis (Petrella et al., 2007). Previously isolated mutations affect both HtsRC and Adducin-related isoforms, resulting in a pleiotropic phenotype involving both germline mitotic divisions as well as ring canal development. To determine the consequence of specifically disrupting HtsRC expression, we used CRISPR/Cas9 non-homologous end joining (NHEJ) mutagenesis to introduce truncation mutations in the exon encoding HtsRC (Figures 1G and 2A, green). Ovaries from these mutant females exhibited none of the germline mitotic defects associated with alleles affecting the Adducin isoforms. The fusome was present and exhibited Adducin localization indistinguishable from wild type (Figure S3). Instead, the *htsRC* alleles displayed a specific ring canal phenotype, in which the robust inner rim of F-actin failed to accumulate (Figures S3, 2F and 2G’). Of note, the *htsRC*-specific actin phenotype can be rescued by expression of Ovhts::GFP (Figure S3). Consistent with previous work, Filamin was detectable at these ring canals (Figure 2G”’), confirming that Filamin localization to ring canals is independent of HtsRC.

The loss-of-function phenotypes of *kelch* and *htsRC* are opposite of one another with respect to the recruitment of a ring canal F-actin cytoskeleton: in *htsRC* truncation mutants, ring canals lack the robust ring of F-actin found in wild-type ring canals (compare Figures 2C’ and 2G’), while in *kelch* mutant females the F-actin cytoskeleton accumulates aberrantly in the ring canal lumens (Figure 2E’). We examined double mutant combinations of loss-of-function alleles *kel*^*DE1*^ and *hts*^*R913fs*^. The double mutant was identical to the *hts*^*R913fs*^ single mutant with regard to the ring canal F-actin (Figures 2F-2I) and oocyte size phenotype (Figure 2J), demonstrating that *htsRC* is epistatic to *kelch*. This is consistent with HtsRC being an important substrate of CRL3^Kelch^: when HtsRC is absent, loss of Kelch has no obvious consequence or additional phenotype.

**Figure 2.**
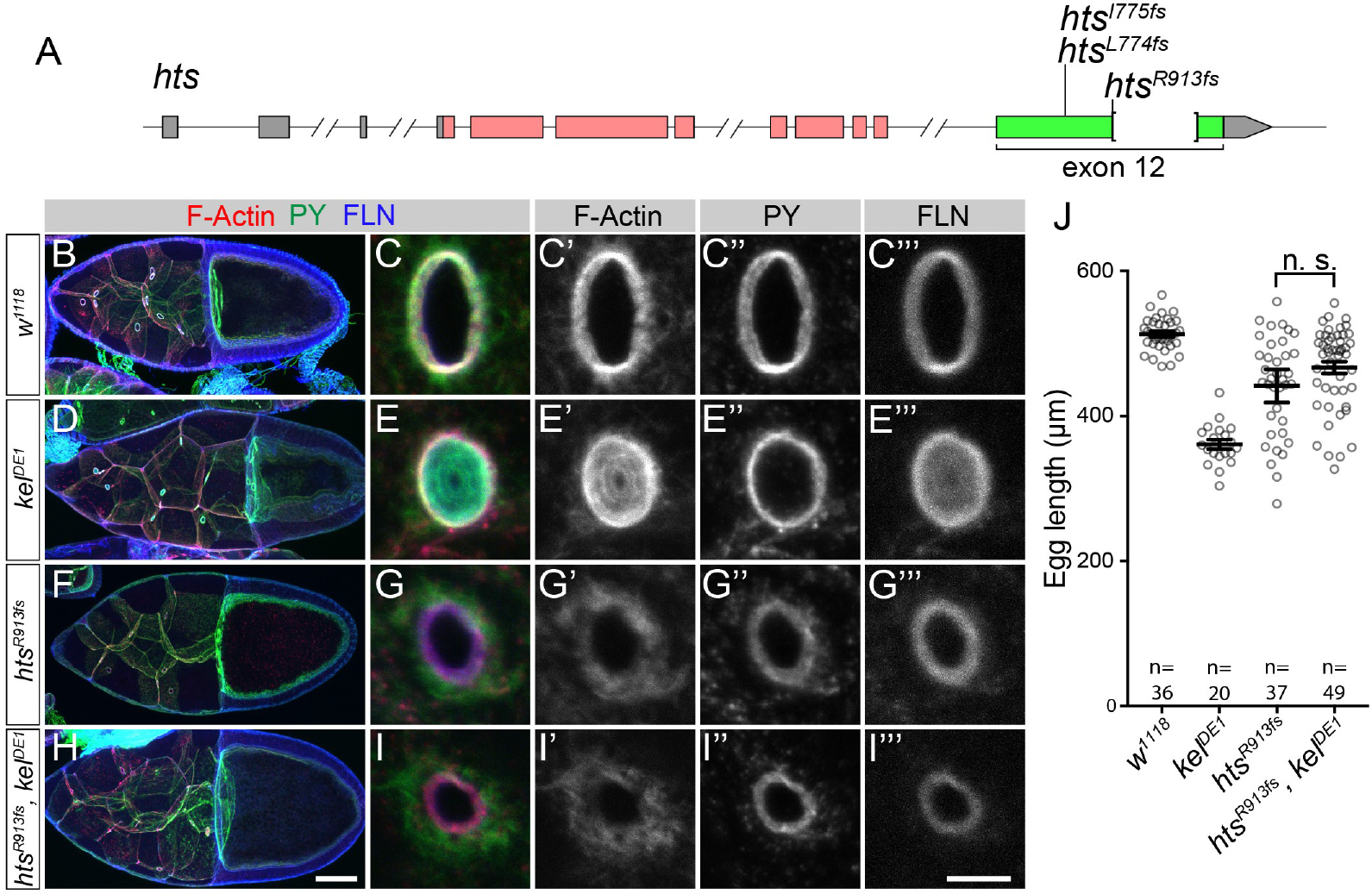
Genetic analysis indicates hts is epistatic to *kelch*. (A) Diagram showing loss-of-function mutations in hts exon 12. (B-I) Wild type, *kel*^*DE1*^, *hts*^*R913fs*^, and *hts*^*R913fs*^, *kel*^*DE1*^ stage 10 egg chambers showing the distributions of F-actin, phosphotyrosine (PY), and Filamin (FLN). kelch mutant egg chambers display a fully penetrant “small oocyte” phenotype at stage 10 (D). Oocytes in *hts*^*R913fs*^ (F) and *hts*^*R913fs*^, *kel*^*DE1*^ double mutant egg chambers (H) do not exhibit a penetrant stage 10 growth defect. (E, G, I) High resolution images of ring canals from the genotypes indicated. *kel*^*DE1*^ and *hts*^*R913fs*^ ring canals exhibit opposite phenotypes with respect to ring canal F-actin accumulation: excess F-actin in *kel*^*DE1*^, and only a small amount of peripheral F-actin accumulation around the ring canals in the *hts*^*R913fs*^ mutant. (J) Quantification of egg lengths from the indicated genotypes. The *hts*^*R913fs*^, *kel*^*DE1*^ double mutant phenotype is indistinguishable from *hts*^*R913fs*^, demonstrating that *hts* is epistatic to *kelch*. Bars represent mean ± 95% confidence interval. One-way ANOVA, Tukey’s multiple comparison test; n.s., not significant. See also Figure S3

### Altered HtsRC expression can suppress or enhance the *kelch*-like ring canal phenotype

To further test the hypothesis that HtsRC is the CRL3^Kelch^ ring canal substrate, we performed a series of genetic enhancement and suppression experiments to determine how HtsRC protein expression levels affect the *kelch*-like ring canal phenotype. First, we tested if HtsRC expression could enhance or suppress the *kelch*-like ring canal phenotype caused by germline-specific proteasome inhibition by RNAi (Figure 3A). Ring canal F-actin thickness was assessed using the full width at half max (FWHM) measure of F-actin intensity plots spanning across ring canals of stage 6 egg chambers. Germline-specific expression of shRNAs targeting ProsBeta5, a catalytic subunit of the proteasome, led to *kelch*-like ring canals (Figure 3A, red), marked by significantly thicker ring canal F-actin compared to wild-type ring canals (Figure 3A, black). Increased HtsRC::GFP expression, achieved by expressing a UASH-Ovhts::GFP construct, significantly enhanced the *kelch*-like ring canal phenotype observed upon proteasome inhibition (Figure 3A, green). (UASH-Ovhts::GFP encodes a full-length Ovhts protein with a C-terminal GFP tag (Petrella et al., 2007).) Additionally, removal of one copy of *kelch* in the context of proteasome inhibition and HtsRC::GFP expression further increased the ring canal F-actin thickness (Figure 3A, yellow). Conversely, removal of one copy of *htsRC* was sufficient to suppress the *kelch*-like ring canal phenotype caused by proteasome inhibition (Figure 3A, blue). These results suggest that increased or decreased HtsRC expression can significantly enhance or suppress, respectively, the *kelch*-like ring canal F-actin phenotype caused by proteasome inhibition and that HtsRC, Kelch, and the proteasome genetically interact.

**Figure 3.**
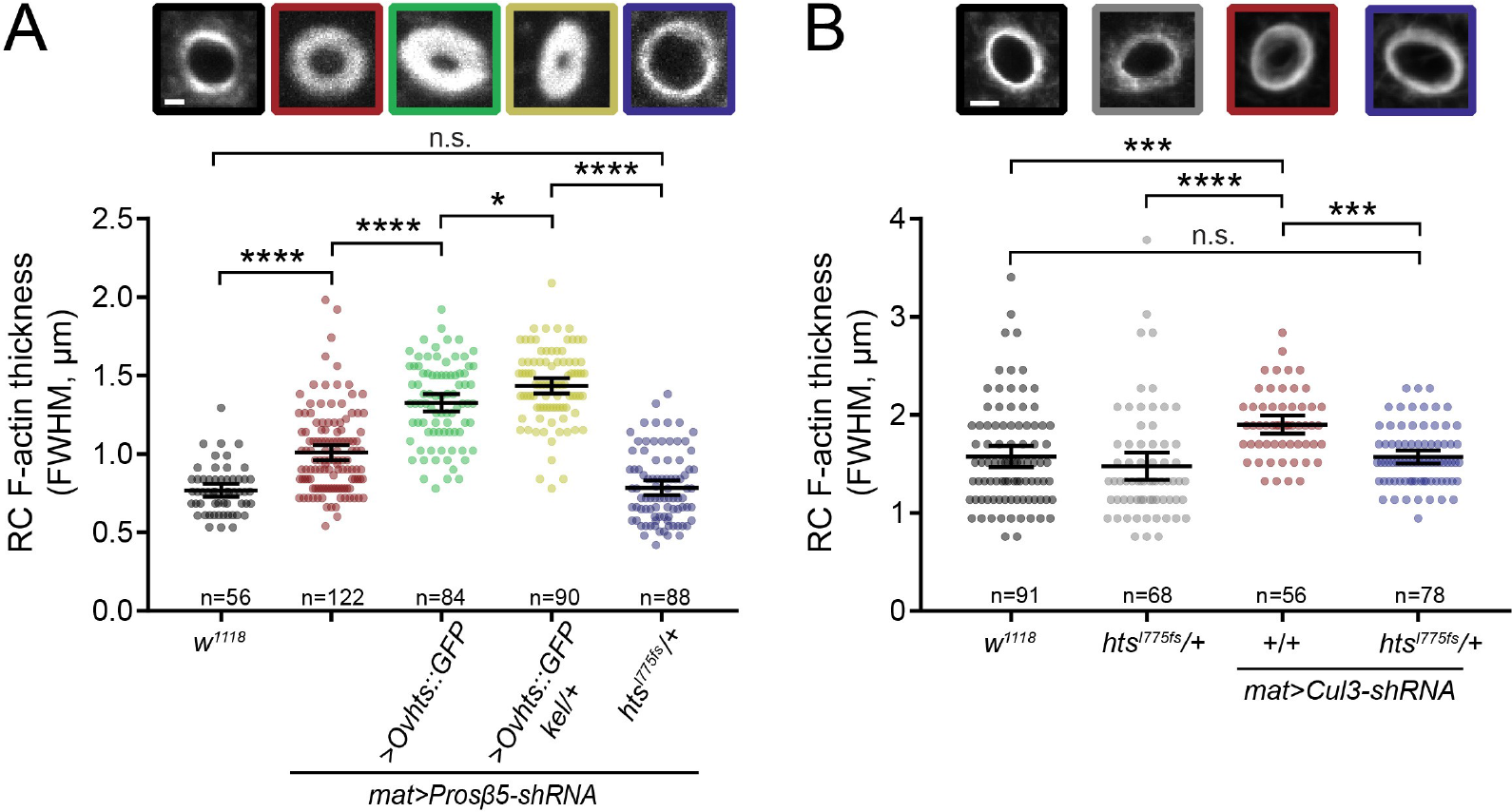
Altered HtsRC expression can suppress or enhance the *kelch*-like ring canal phenotype. (A-B) Ring canal F-actin thickness in stage 6 (A) or stage 9 (B) egg chambers was calculated using the full width at half max (FWHM) measure of F-actin intensity plots spanning across ring canals. Colored boxed images show representative ring canals of each genotype, with data points below for each ring canal measured. Bars represent mean F-actin thickness ± 95% confidence interval. n denotes number of ring canals measured. (A) Increased or decreased HtsRC expression enhances or suppresses, respectively, the *kelch*-like ring canal phenotype observed upon germline-specific proteasome inhibition by RNAi. Proteasome inhibition leads to *kelch*-like ring canals, marked by significantly thicker F-actin rings (red). Increased HtsRC::GFP expression (green) and increased HtsRC::GFP expression with loss of one copy of *kelch* (yellow) significantly increases ring canal F-actin thickness, whereas removal of one copy of *htsRC* fully suppresses the *kelch*-like phenotype observed upon proteasome inhibition (blue). (B) Germline-specific knockdown of *Cul3* by RNAi leads to *kelch*-like ring canals (red), marked by significantly thicker F-actin rings. Removal of one copy of *htsRC* is sufficient to suppress the *kelch*-like phenotype (blue). (****) p<0.0001, (***) p<0.001; (*) p<0.05; one-way ANOVA, Tukey’s multiple comparison test. n.s., not significant.

To test the ability of HtsRC expression to affect the ring canal F-actin organization in a different context, we analyzed ring canals in stage 9 control egg chambers or egg chambers experiencing germline-specific knockdown of *Cul3* (Figure 3B). As shown previously (Hudson et al., 2015), knockdown of *Cul3* leads to *kelch*-like ring canals (Figure 3B, red). Removal of one copy of *htsRC* was sufficient to fully rescue this phenotype (Figure 3B, blue), as the ring canal F-actin thickness was restored to wild-type levels. These data show that alteration of HtsRC protein levels can enhance and suppress the *kelch*-like ring canal F-actin phenotype in multiple contexts.

### HtsRC ring canal protein levels are dependent on Kelch

If HtsRC levels at ring canals are regulated by CRL3^Kelch^-mediated ubiquitylation, we would expect partial loss-of-function mutations in *hts* may be suppressed by mutations in *kelch*. We reduced the expression of HtsRC using an shRNA construct targeting *hts* driven by *matGal4*. The *matGal4* driver expresses Gal4::VP16 from the maternal *αTub67C* promoter, and results in strong expression beginning late in germarial development (Figure S5), after the earlier requirement for *hts* in the germline mitotic divisions. This results in a strong reduction, but not elimination of the HtsRC protein (Yan et al., 2014) (Figures 4A and 4B). When *hts* shRNA is expressed in a *kelch* mutant background, HtsRC levels at ring canals are increased compared to expression of *hts* shRNA in wild type (Figure 4C).

If HtsRC is indeed a CRL3^Kelch^ substrate, then its protein levels should be dependent on Kelch. To test this assumption, we tested the effect on HtsRC::GFP levels of over-expressing Kelch (Figures 4D and 4E). HtsRC::GFP fluorescent protein levels were drastically reduced in egg chambers co-expressing mCherry::Kelch compared to mCherry control (Figures 4D and 4E). Furthermore, western analysis revealed that total HtsRC protein species decreased significantly upon co-expression of mCherry::Kelch compared to mCherry control (Figure S4). Of note, the levels of HtsRC::GFP species were most dramatically decreased upon Kelch overexpression, while endogenous HtsRC remained mostly unchanged (Figure S4).

**Figure 4.**
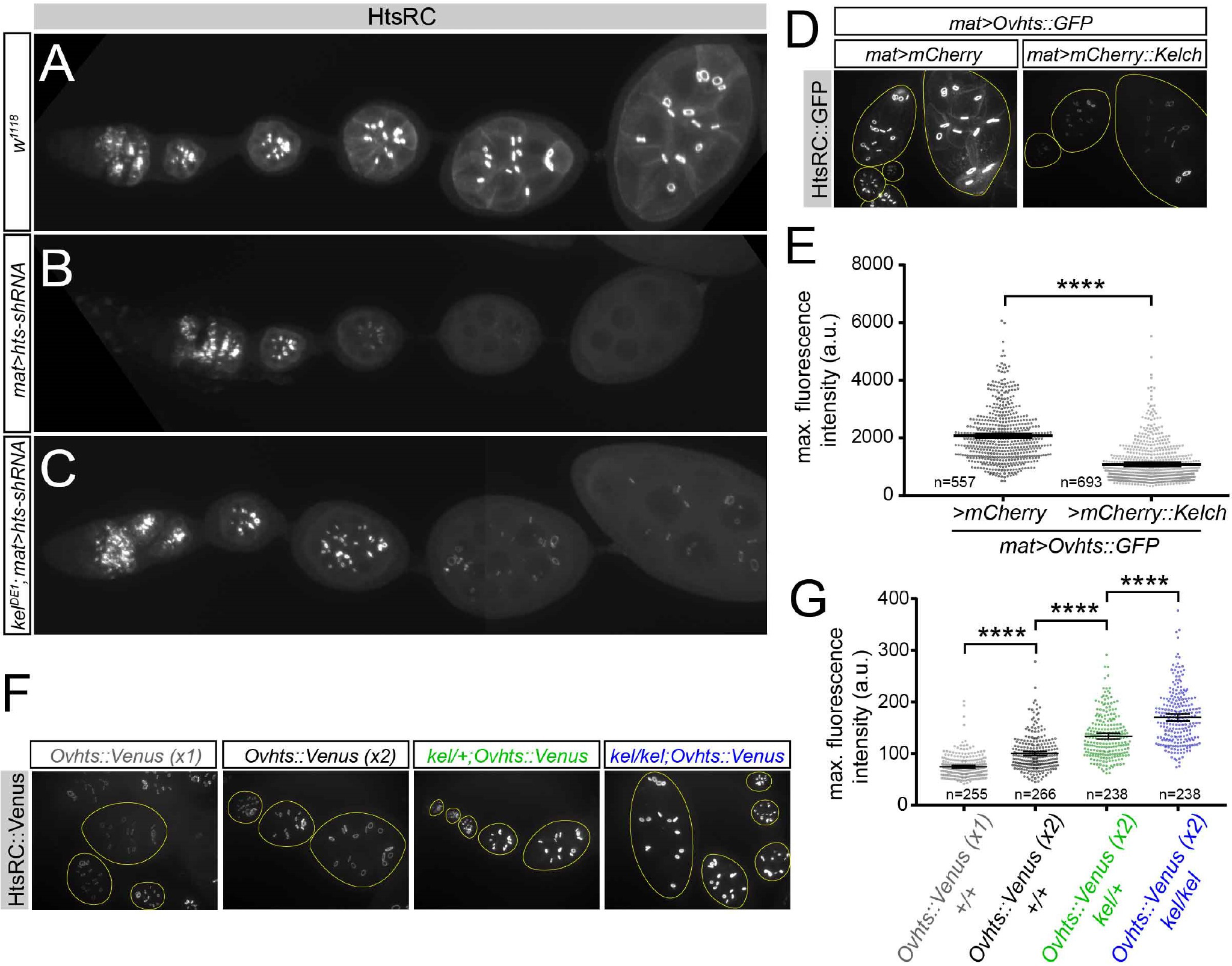
HtsRC ring canal protein levels are dependent on Kelch. (A) Wild-type HtsRC expression in the germarium and subsequent stages. (B) *matGal4*-driven expression of a shRNA against *hts* results in near elimination of HtsRC from the germline in stages where *matGal4* expression is high. (C) Mutations in *kelch* suppress the *hts* shRNA knockdown phenotype, resulting in detectable HtsRC in stage 2-8 egg chambers. (D) HtsRC::GFP levels decrease as Kelch protein levels increase. HtsRC::GFP fluorescent protein levels decrease with mCherry::Kelch co-expression compared to mCherry control expression. All constructs are driven by *matGal4*. Yellow circles outline individual egg chambers. (E) Quantification of HtsRC::GFP fluorescent protein levels at ring canals. (****) p<0.0001; Student’s t-test.(F) HtsRC::Venus fluorescence levels at ring canals increase in a dose-dependent manner depending on *kelch* gene copy levels. Yellow circles outline individual egg chambers. (G) Quantification of HtsRC::Venus fluorescent protein levels at ring canals. The mean maximum fluorescence intensity of the *Ovhts::Venus x2* sample (black) was normalized to 100 relative fluorescence units. For E and G, data points in the scatter plot represent the maximum fluorescence intensity of HtsRC::Venus or HtsRC::GFP for each ring canal measured. Bars represent mean maximum fluorescence intensity of HtsRC::Venus or HtsRC::GFP ± 95% confidence interval. n denotes number of ring canals analyzed. (****) p<0.0001; one-way ANOVA, Tukey’s multiple comparison test. See also Figures S4 and S5.

Finally, we examined the levels at ring canals of HtsRC::Venus fluorescent fusion protein expressed from a BAC transgene in the context of varying doses of *kelch* gene copies (Figures 4F and 4G). Fixed egg chambers were imaged with a constant exposure time, and maximum intensity projections were rendered in FIJI software to visualize HtsRC::Venus levels (Figure 4F). HtsRC::Venus fluorescent protein levels progressively increased as *kelch* gene copy levels decreased (Figure 4F). This effect was quantified by calculating the maximum fluorescence intensity of HtsRC::Venus for individual ring canals analyzed in four genotypes (Figure 4G). The mean maximum fluorescence intensity of HtsRC::Venus progressively increased as *kelch* gene copy was reduced (compare black, green, blue data). These data show that Kelch protein levels directly affect ring canal-localized HtsRC protein levels.

### Removal of the Kelch NTR results in hyper-active Kelch

Kelch contains an N-terminal region (NTR) of 120 amino acids that is rich in low-complexity sequence (Figure 5A). We previously analyzed a transgene encoding a Kelch protein lacking the NTR expressed at low levels using the *otu* promoter (Robinson and Cooley, 1997). *otu*-driven expression of Kelch^ΔNTR^ resulted in a dominant-negative female-sterile phenotype due to the destabilization of germ cell membranes. This phenotype was attributed to the premature localization of Kelch^ΔNTR^ to ring canals, as the Kelch^ΔNTR^ protein localized earlier in oogenesis than wild-type Kelch expressed from control constructs (Robinson and Cooley, 1997). Given our results indicating that Kelch regulates HtsRC levels as an E3 ligase in the UPS, we reexamined the phenotype caused by Kelch^ΔNTR^ expression. We created a UASp-Kelch^ΔNTR^ transgene to express the ΔNTR protein with greater control over the timing and level of expression. When we expressed either wild-type Kelch or Kelch^ΔNTR^ at high levels using *matGal4*, the females expressing wild-type Kelch were fertile and displayed normal localization of HtsRC (Figure 5B). In contrast, the females expressing Kelch^ΔNTR^ were sterile, similar to what we had observed previously with *otu*-Kelch^ΔNTR^ constructs. Immunofluorescence analysis of HtsRC in the Kelch^ΔNTR^-expressing egg chambers revealed that HtsRC was nearly undetectable within the expression domain of *matGal4* (Figure 5C, C’), and *mat>Kelch*^Δ*NTR*^ egg chambers failed to complete oogenesis.

Expression of Kelch^ΔNTR^ at low levels using the weak *otu-Gal4::VP16* driver (*otuGal4*) (Figure S5) resulted in a similar but less severe phenotype, and western blotting revealed reduced levels of HtsRC (Figure 5D and E). When we compared the expression levels of Kelch^ΔNTR^ with wild-type Kelch driven by *otuGal4*, we found that Kelch^ΔNTR^ was expressed at much higher levels than wild-type Kelch, suggesting that loss of the NTR resulted in a stabilized protein (unpublished). This raised the possibility that elevated CRL3^Kelch^ activity toward HtsRC may have resulted from the increased quantities of a stabilized Kelch^ΔNTR^. This would be similar to human mutations in KLHL24 that result in the production of a BBK protein that lacks its N-terminal region, leading to increased levels of KLHL24 and elevated levels of CRL3^KLHL24^ activity (Lin et al., 2016). To address the effect of differing levels of Kelch, we compared the effects of Kelch^ΔNTR^ driven by *otuGal4* with wild-type Kelch driven by *matGal4* (see Figure S5 for Gal4 expression comparisons). Wild-type Kelch driven by *matGal4* was present at twice the level of Kelch^ΔNTR^ driven by *otuGal4* (Figure 5D and F); however, HtsRC levels were only reduced upon expression of Kelch^ΔNTR^. This suggests that altered regulation of Kelch^ΔNTR^, rather than increased quantity, is responsible for the apparent gain-of-function effects seen with Kelch^ΔNTR^ expression. Removal of the NTR appears to result in a hyper-active form of Kelch.

### A PEAEQ sequence motif in HtsRC is necessary for its interaction with Kelch

Identification of the conserved PEAEQ sequence in HtsRC and other KREP-interacting proteins suggests that this motif is a binding site for the Kelch KREP domain. If this is the case, and if HtsRC ubiquitylation is essential for ring canal cytoskeletal organization, then mutations in the *hts* PEAEQ motif that inhibit Kelch binding should result in a *kelch*-like phenotype. To test this, we used CRISPR/Cas9 NHEJ mutagenesis to create mutations in the PEAEQ sequence. sgRNAs targeting codons for two conserved residues in the PEAEQ sequence, E922 and Q925, were used to generate germline mutations (Figure 6A).

**Figure 5.**
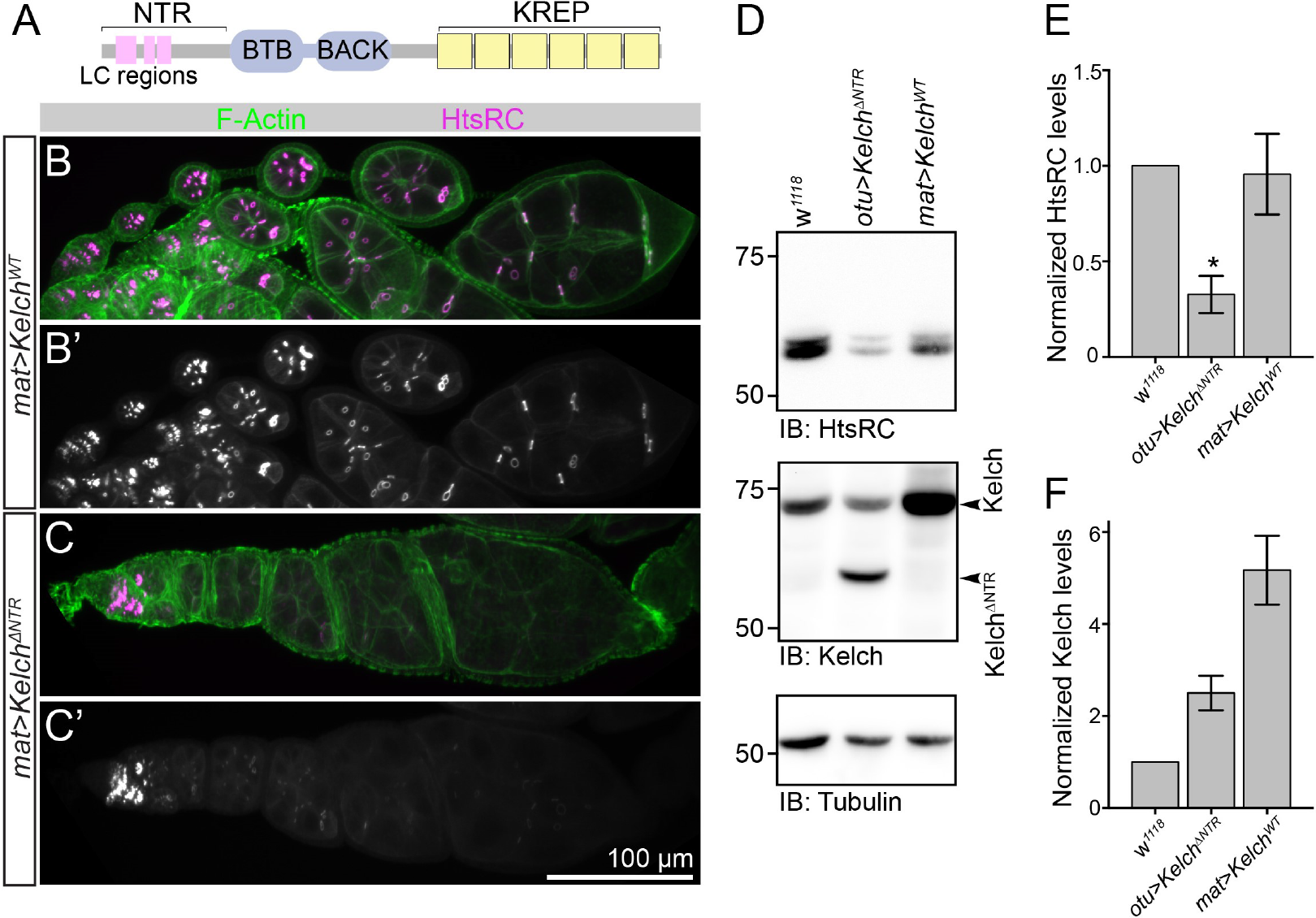
Removal of the Kelch NTR results in hyper-active Kelch. (A) Cartoon of Kelch protein motifs, with low-complexity regions (LC) in the NTR (pink boxes). (B, B’) Expression of wild-type protein with the *matGal4* driver results in a normal distribution of HtsRC in egg chambers. (C, C’) Expression of Kelch^ΔNTR^ using the *matGal4* driver results in dominant phenotypes consistent with Kelch^ΔNTR^ acting as a hyperactive mutant. *matGal4*-driven expression of Kelch^ΔNTR^ in the vitellarium results in dramatic loss of HtsRC fluorescence compared to expression of wild-type Kelch. Note that HtsRC levels in wild type and *matGal4>Kelch*^Δ*NTR*^ are comparable in the germarium, where the *matGal4* driver is not expressed. (D and E) Western analysis of HtsRC and Kelch levels. Expression of Kelch^ΔNTR^ results in reduced HtsRC protein compared to *w*^*1118*^ or *matGal4>Kelch*^*WT*^. (*) p<0.05. (F) Kelch^ΔNTR^ expression driven by *otuGal4* results in approximately twice the level of endogenous Kelch. *matGal4*-driven expression of wild-type Kelch results in an approximately 5-fold increase in Kelch expression. See also Figure S5.

We recovered in-frame deletions of both E922 and Q925 in the PEAEQ motif, and in marked contrast to truncation mutations that lack a robust ring canal F-actin cytoskeleton (Figures 2 and S2), the in-frame deletions resulted in phenotypes similar to *kelch* loss-of-function mutants. The stronger allele, a deletion of the E922 codon (*hts*^Δ*E922*^), resulted in a phenotype indistinguishable from the *kel*^*DE1*^ null allele (Figures 6B-6D): F-actin and the mutant HtsRC protein accumulated in ring canals, and the flies were completely sterile. The levels of HtsRC at ring canals were 2-fold higher in *hts*^Δ*E922*^ and *kel*^*DE1*^ mutants compared to wild type (Figure 6E). In addition, we found that in *hts*^Δ*E922*^ mutants, Kelch no longer localized to ring canals (Figures 6G-6I), despite Kelch protein being present in *hts*^Δ*E922*^ mutants (Figure 6F). Curiously, while increased levels of HtsRC at ring canals in *kelch* or *htsRC* PEAEQ mutants were obvious by immunofluorescence, we detected no difference in their steady-state accumulation by western blot, (Figure 6H). The *hts*^Δ*Q925*^ allele appeared to be weaker, and levels of HtsRC measured at ring canals were intermediate between wild type and *hts*^Δ*E922*^ mutants (Figures S6A-S6C and S6G). Consistent with this result, Kelch was detected at *hts*^Δ*Q925*^ ring canals, but at reduced levels compared to wild type (Figures S6D-S6F and S6H). Together, these results provide compelling in vivo evidence that the PEAEQ sequence is a binding site for Kelch.

**Figure 6.**
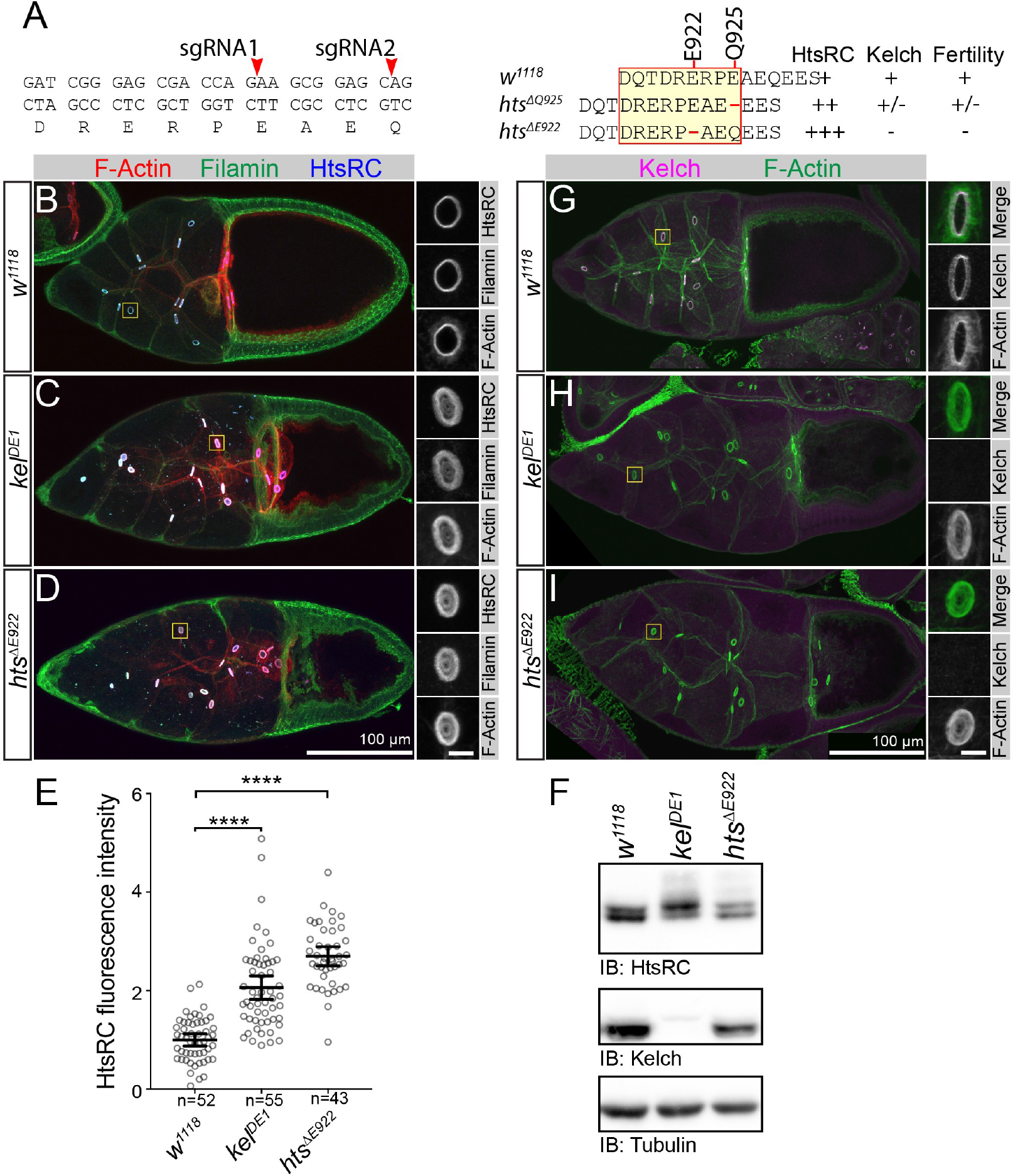
A PEAEQ sequence motif in HtsRC is necessary for its interaction with Kelch. (A) Location of sgRNAs targeting the HtsRC PEADQ motif (left), and summary of single-codon deletion mutations in PEAEQ sequence (right). Red arrowheads indicate locations where sgRNAs direct Cas9-mediated cleavage. *hts*^Δ*E922*^ is an in-frame deletion resulting in a single amino acid deletion of the conserved E922 residue; *hts*^Δ*Q925*^ deletes Q925. (B-D) Wild type, *hts*^Δ*E922*^, and *kelch*^*DE1*^ egg chambers labeled to reveal F-actin, HtsRC, and Filamin. Homozygous *hts*^Δ*E922*^ egg chambers and ring canals are indistinguishable from *kel*^*DE1*^. (E) Measured fluorescence intensity of HtsRC immunolabeled ring canals. Bars represent mean ± 95% confidence interval. (****) p<0.0001; one-way ANOVA with Dunnett’s multiple comparison test. (G-I) Kelch localization in WT, *hts*^Δ*E922*^, and *kelch*^*DE1*^ egg chambers. Kelch localizes to ring canals in wild-type egg chambers (G), but not in *hts*^Δ*E922*^ mutants (I), similar to *kelch* loss of function mutations (H), despite being expressed at normal levels (F). For B-D and G-I, ring canals within yellow box are shown in inset to the right, with scale bar equal to 5 *μ*m. (F) Western blot showing steady-state levels of HtsRC and Kelch; Tubulin serves as a loading control. Overall HtsRC levels are not significantly increased compared to wild type. See also Figure S6.

### HtsRC is ubiquitylated and degraded by the proteasome

If HtsRC is a CRL3^Kelch^ substrate, then it should be subjected to ubiquitylation. To assess HtsRC ubiquitylation, we expressed HtsRC in cultured S2 cells by cloning exon 12 (see Figure 1G) into an expression vector with HA epitope tags at both the N- and C-termini (Figure 7A). We reasoned that transfected HtsRC — despite it not being endogenously expressed in these cells — could still be subjected to its normal mode of regulation by CRL3^Kelch^ since all components of the CRL3^Kelch^ ubiquitin ligase machinery are present. S2 cells produced both uncleaved 3xHA::HtsRC::HA precursor protein and the HtsRC::HA cleavage product with no N-terminal tag (Figure 7A). 24 hours post-transfection, cells were treated with DMSO (control) or 1 *μ*M Bortezomib, a proteasome inhibitor, for 3 and 6 hours. Whole cell lysates analyzed by western blotting showed that proteasome inhibition resulted in a higher molecular weight, presumably ubiquitylated, HtsRC species (Figure 7B). Proteasome inhibition also resulted in a two-fold increase of HtsRC protein compared to DMSO control (Figures 7C and 7D).

**Figure 7.**
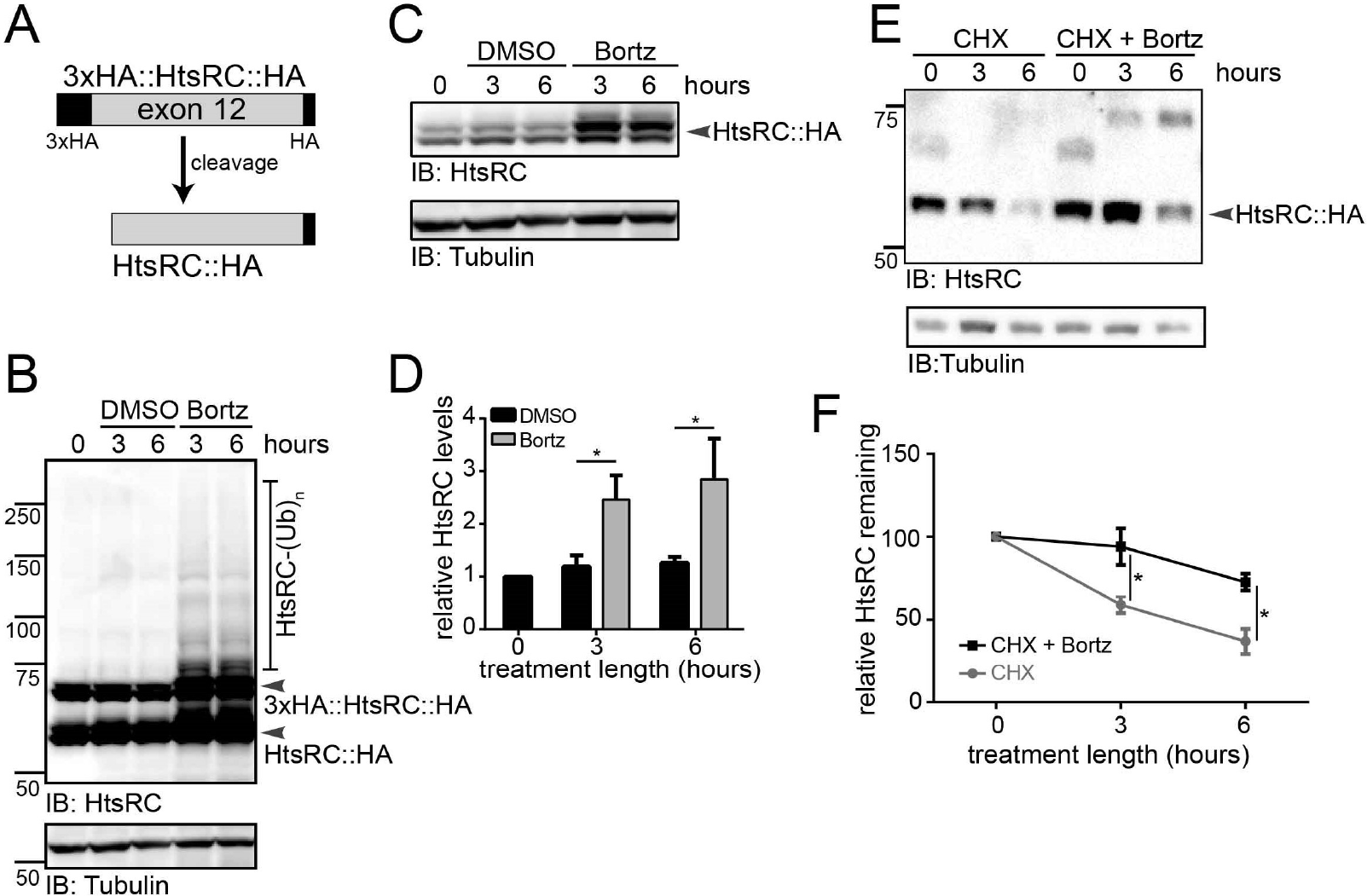
HtsRC is uniquitylated and degraded by the proteasome. (A) Cartoon showing full-length 3xHA::HtsRC::HA and cleaved HtsRC::HA protein products. (B) S2 cells expressing 3xHA::HtsRC::HA driven by the actin promoter were treated with DMSO or 1 *μ*M Bortezomib for 3 or 6 hours, 24 hours post-transfection. Cells were harvested and cell lysates were analyzed by western analysis. Proteasome inhibition results in higher molecular weight, presumably ubiquitylated, HtsRC species. (C-D) S2 cells were treated with control DMSO or 1 *μ*M Bortezomib for 3 and 6 hours, 36 hours post-transfection. Cell lysates were analyzed by western analysis. Levels of HtsRC::HA were elevated after Bortezomib treatment. (*) p<0.05; Student’s t-test. Data are from three independent experiments. (E) 24 hours post-transfection of 3xHA::HtsRC::HA, S2 cells were treated with 100 *μ*g/mL Cycloheximide (CHX) with or without 1 *μ*M Bortezomib, and cells were harvested at 3 and 6 hour timepoints for western analysis. (F) Quantification of HtsRC protein levels by western analysis of three independent CHX chase experiments. HtsRC protein levels decrease after CHX treatment and are stabilized with CHX and Bortezomib treatment, suggesting that HtsRC is degraded by the proteasome. (*) p<0.05; Student’s t-test.

We next tested HtsRC stability upon proteasome inhibition in the context of inhibiting new protein synthesis with cycloheximide (Figures 7E and 7F). In cycloheximide-treated S2 cells, HtsRC levels declined over 6 hours. With the addition of proteasome inhibition, HtsRC declined more slowly (Figures 7E and 7F). These data suggest that HtsRC is degraded by the proteasome in cultured Drosophila cells.

## Discussion

In this study, we show that the CRL3^Kelch^ targets HtsRC at ring canals for ubiquitylation and destruction, providing a mechanism for how Kelch shapes the cytoskeleton during ring canal growth. We provide four lines of evidence showing that HtsRC is the key substrate of CRL3^Kelch^: first, we detected HtsRC in a complex with the substrate-binding KREP domain of Kelch; second, we showed that levels of HtsRC are sensitive to the dosage of Kelch; third, we identified a putative binding motif in HtsRC that, when mutated, recapitulates the *kelch* phenotype and results in a loss of Kelch localization to RCs; and finally, we show that HtsRC can be ubiquitylated in cultured cells. Together, these results provide compelling evidence that HtsRC levels are controlled by CRL3^Kelch^-mediated targeted destruction to promote the coordinated expansion of the ring canal cytoskeleton.

A striking feature of many proteins identified by mass spectrometry in complex with the KREP domain was the presence of a linear PEAEQ peptide motif. Two human BBK proteins, KLHL2 and KLHL3, are most closely related to Drosophila Kelch, and both have been shown to bind a similar EPEEPEADQ motif in WNK4, a substrate of KLHL3 (Schumacher et al., 2014). A structure of a KLHL3ȓWNK4 peptide complex revealed that the EPEEPEADQ peptide binds on the top surface of the KLHL3 KREP *β*-propeller, making extensive contacts with inter-strand surface loops derived from blades 2-4. We propose that the HtsRC PEAEQ motif binds Drosophila Kelch in a similar manner, and the prevalence of PEAEQ proteins we identified by both pulldown and yeast two-hybrid screening suggests that proteins with PEAEQ-like linear motifs are potential targets of Kelch and closely-related BBK proteins.

In contrast, Keap1, a more distantly related BBK protein, targets its substrate Nrf2 for ubiquitylation through a high-affinity interaction with a short DxETGE sequence (Lo et al., 2006). A structure of the Keap1ȓDxETGE peptide complex revealed that Keap1 also engages its target peptide through contacts with surface loops extending from the top of the Keap1 *β*-propeller structure, similar to the KLHL3ȓPEADQ interaction. Unlike KLHL2 and 3 however, the Nrf2 DxETGE peptide was found to be closely associated with surface loops derived from blades 1, 5 and 6; it therefore appears that the Keap1 KREP domain binds the DxETGE peptide in a manner distinct from the KLHL3ȓPEADQ interaction. Based on these observations, we speculate that subfamilies of BBK proteins may engage substrates through distinct short peptide motifs.

Interactions between ubiquitin ligases and their substrates are typically regulated such that substrate ubiquitylation is restricted to the appropriate time and place. Several CRLs involved in germline development illustrate a number of unusual mechanisms. The Germ cell-less protein (GCL) is a component of a CRL3 that degrades the Torso receptor tyrosine kinase in order to maintain primordial germ cell fate (Pae et al., 2017). GCL is located at the nuclear envelope during interphase, and only free to act following nuclear envelope breakdown during mitosis. Mutants that disrupt this mode of regulation result in the apparent ubiquitylation of inappropriate substrates and severe germ cell defect (Pae et al., 2017). KLHL10, a BBK CRL3 substrate adaptor required for spermatid differentiation, is regulated by Scotti, a protein that acts as a pseudosubstrate inhibitor of CRL3^KLHL10^ to regulate its activity in a graded fashion during spermatid elongation (Arama et al., 2007; Kaplan et al., 2010).

Developmental regulation of ubiquitin ligaseȓsubstrate interactions often involves phosphorylation; in Drosophila oogenesis, an isoform of the RNA-binding protein Oskar is targeted for degradation by SCF^Slimb^ upon phosphorylation (Morais-de Sa et al., 2013). We note that HtsRC migrates as a doublet on SDS-PAGE gels (Figure 6F), possibly indicating the presence of phosphorylated forms; determining whether substrate phosphorylation regulates HtsRC degradation will be an area of future investigation. Phosphorylation of the E3 ligase can also regulate substrate ubiquitylation; KLHL3, the human BBK substrate adaptor that targets WNK4, is phosphorylated on a serine residue shown to make contacts with the WNK4 PEADQ motif (Shibata et al., 2014). This serine is conserved in Drosophila Kelch (S535 in Kelch), and our MS/MS analysis of the purified KREP domain detected the presence of phosphorylated S535 (Hudson, unpublished). However, mutating S535 to either a non-phosphorylatable or phosphomimetic form did not affect the ability of either mutant protein to rescue the *kelch* mutant ring canal phenotype (Hudson, unpublished), indicating that phosphorylation of this serine is not essential for targeting of HtsRC by Kelch.

We did find evidence that the Kelch N-terminal region is important in regulating CRL3^Kelch^ activity. We previously found that transgenic constructs driving low-level germline expression of Kelch^ΔNTR^ resulted in a dominant female-sterile phenotype (Robinson and Cooley, 1997). This phenotype was thought to arise from premature localization of Kelch to ring canals, resulting in later membrane destabilization. Our current results demonstrating that Kelch^ΔNTR^ expression depletes HtsRC from germline cells suggests that the phenotype of Kelch^ΔNTR^ results instead from a hyperactive CRL3^Kelch^ protein that targets HtsRC in an unregulated manner. We note that the phenotype of Kelch^ΔNTR^ is similar to, but more severe than loss-of-function mutations in *htsRC*, suggesting that the Kelch^ΔNTR^ may target proteins that are not normally substrates of CRL3^Kelch^ during oogenesis.

The Drosophila Kelch NTR is a 120 amino-acid extension rich in low-complexity sequence. It is conserved in Kelch orthologs in insects, but not in KLHL2 and KLHL3. However, 13 human BBK proteins have N-terminal extensions greater than 60 amino acids, and many of these are rich in low-complexity sequence. Interestingly, gain-of-function mutations in the one of these genes, KLHL24, produce a mutant protein lacking part of its NTR. These mutant KLHL24 proteins are stabilized and accumulate to high levels, resulting in unregulated ubiquitylation and degradation of its substrate, Keratin 14, which causes skin fragility (Lin et al., 2016). Stabilization of mutant KLHL24 results from impaired auto-catalytic ubiquitylation and degradation of KLHL24, a commonly observed feature or CRLs (de Bie and Ciechanover, 2011). While Kelch also undergoes Cul3-dependent autocatalytic degradation (Hudson and Cooley, 2010; Hudson et al., 2015), we do not think the hyperactive phenotype of Kelch^ΔNTR^ is simply a consequence of increased protein levels since elevated levels of wild-type Kelch do not have a similar phenotype.

**Figure 8.**
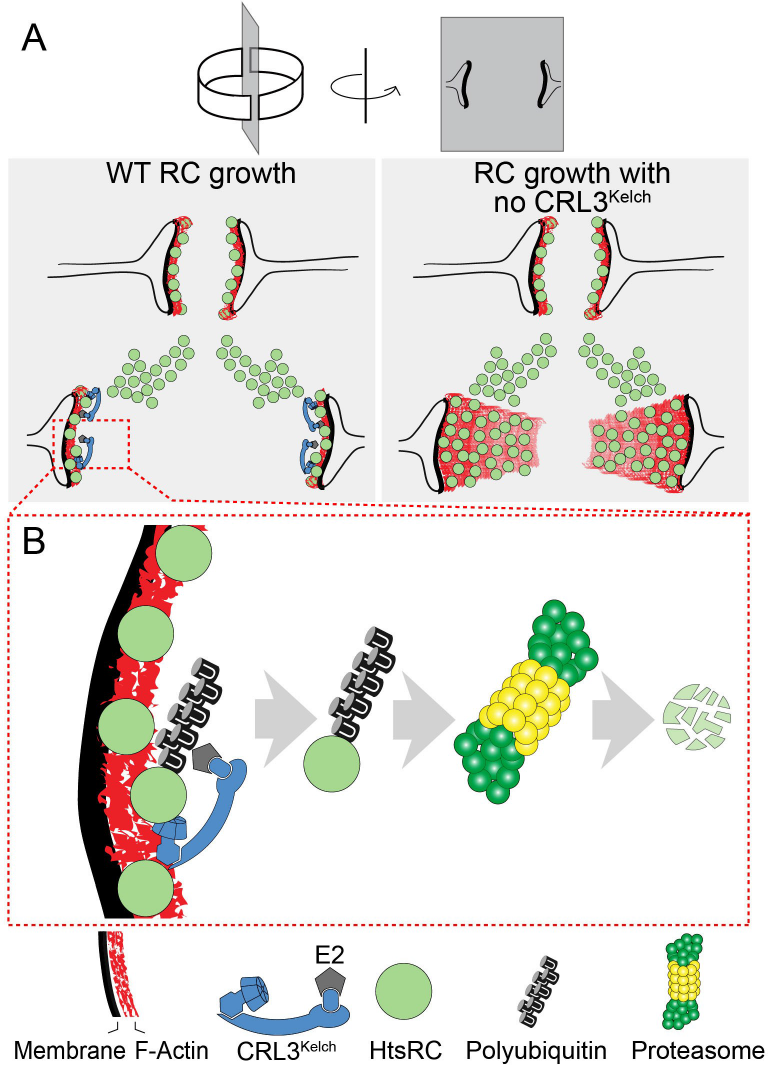
Model for ring canal growth regulated by CRL3^Kelch^. (A) Wild type and *kelch* ring canals diagrammed in cross section. Ring canals recruit HtsRC, which results in the accumulation of a robust F-actin cytoskeleton that supports ring canal growth. (B) In wild type, CRL3^Kelch^ ubiquitylates HtsRC, targeting it for destruction by the proteasome. Removal of HtsRC by CRL3^Kelch^-mediated destruction is required for the disassembly of the ring canal cytoskeleton from the ring canal lumen as the ring canal expands.

Based on our results, we propose that HtsRC is the key driver of ring canal growth and that its levels are regulated by the action of CRL3^Kelch^ and the ubiquitin-proteasome system (Figure 8). HtsRC is essential for the dynamic and robust F-actin cytoskeleton that lines the inner membrane of the ring canal; F-actin fails to accumulate in *hts* mutants (Petrella et al. (2007); Yue and Spradling (1992); this study) and F-actin can be restored in these mutants following expression of Ovhts::GFP (Petrella et al., 2007). These results support a model where production and localization of HtsRC mediates ring canal F-actin recruitment and ring canal growth. The molecular mechanism by which HtsRC recruits F-actin to ring canals is not known; sequence analysis of HtsRC reveals no recognizable motifs aside from a predicted coiled-coil structure at its C-terminus. One notable biochemical feature of HtsRC is its poor solubility. Ovarian lysates contain little soluble HtsRC, and our attempts to produce soluble recombinant protein in its native state for in vitro studies have had limited success (unpublished). We speculate that the HtsRC at the ring canal may function by forming an insoluble matrix on which the dynamic ring canal F-actin cytoskeleton is assembled and drives the expansion of the ring canal diameter. However, to allow for the coordinated expansion of the ring canal lumen, HtsRC and associated proteins must be disassembled at the luminal surface, and this requires targeting by CRL3^Kelch^ and the UPS. This represents an unusual mechanism for cytoskeletal remodeling, which generally relies on non-destructive mechanisms to create and dismantle cytoskeletal assemblies. However, if HtsRC functions in a highly insoluble form, removal by the UPS may be the only effective way to control its activity.

## Materials and Methods

### Contact for Reagent and Resource Sharing

Further information and requests for resources and reagents should be directed to and will be fulfilled by the Lead Contact, Lynn Cooley.

### Experimental Model and Subject Details

#### Drosophila genetics

Drosophila were maintained at 25°C on standard fly food medium. Prior to ovary dissections, females were fattened on wet yeast paste overnight at 25°C. See Reagents Table for a detailed list of fly stocks used in this study.

#### S2 cell culture

Drosophila S2 cells (DGRC) were cultured at 27°C in Schneider’s Drosophila Media (Gibco) supplemented with 10% fetal bovine serum and 1x antibiotic-antimycotic (Gibco).

### Method Details

#### Affinity purification and mass spectrometry

The *cher*^1^allele (89E) was recombined with the *UASp-TAPmCherry* and *UASp-TAPmCherry::KREP* constructs integrated at the attP2 site (68A4), and also recombined with the *matGal4::VP16* (mapped to 91D4 by inverse PCR). Recombinant chromosomes were balanced with a TM3, hs-hid balancer (Bloomington #1558) to allow for heat-shock induced selection of desired cross progeny. Approximately 500 ovaries from *cher*^1^ mutant females expressing either TAPmCherry or TAPmCherry::KREP were homogenized in SBP lysis buffer (50 mM HEPES pH 7.5, 150 mM NaCl, 2 mm EDTA, 0.5%Triton X-100, 10% glycerol, 1 mM PMSF, and 5 *μ*g/mL each chymostatin, leupeptin, antipain, pepstatin) in a glass Duall homogenizer with a teflon pestle driven by motorized spinning. Lysates were clarified by centrifugation for 30’ at 20k × g at 4°C, and approximately 40 mg of clarified supernatants were incubated with 75 *μ*L streptavidin beads (Thermo Fisher Scientific #53114). Beads were washed 2x in SBP lysis buffer followed by 2 additional washes in His-tag buffer (50 mM NaPi pH 8, 300 mM NaCl, 5 mM imidazole, 0.2% Triton X-100). Proteins bound to streptavidin beads were eluted in 1 mL 10 mM biotin in His-tag buffer for 20 min at 4°C. Eluted proteins were incubated with 75 *μ*L 50% Ni^++^-NTA slurry for 2 hours, washed 3x in His-tag buffer, and eluted 2x in 35 *μ*L SDS-PAGE sample buffer containing 200 mM DTT and 200 mM imidazole. 25 *μ*L of each sample was loaded on a 4-12% gradient NuPAGE gel and stained with EZBlue colloidal coomassie (Sigma-Aldrich). A second 25 *μ*L aliquot was analyzed at MS Bioworks (Ann Arbor, MI) by running the samples on a 10% Bis-Tris NuPAGE gel, slicing each lane into 10 equal fragments, and subjecting each fragment to in-gel trypsin digestion followed by LC-MS/MS on a Thermo Fisher Q Exactive instrument. MS data were searched against the Uniprot Drosophila proteome at MS Bioworks using Mascot (Matrix Science) and the results were parsed into Scaffold (Proteome Software) for further analysis.

#### Plasmids

To make pAHW-HtsRC::HA for expression of HtsRC in S2 cells, exon 12 of HtsRC with a C-terminal 1xHA tag was PCR-amplified to contain flanking attB1 and attB2 sites. The PCR product was recombined into the Gateway Donor vector pDONR201 in a Gateway BP Clonase II reaction (Thermo Fisher Scientific). The entry clone was further recombined into the Gateway Expression vector pAHW (Drosophila Gateway Vector Collection; Drosophila Genomics Resource Center #1095) in an LR Clonase II reaction Thermo Fisher Scientific). The resulting pAHW-HtsRC::HA expression plasmid allows for expression of HtsRC exon 12 with a 3xHA N-terminal tag and 1xHA C-terminal tag, driven by the Actin5C promoter. pUASp-Kelch^ΔNTR^ was generated by PCR amplifying a region of the Kelch CDS encoding amino acids Q121ȓM689, the same range used in earlier ΔNTR constructs (Robinson and Cooley, 1997). PCR primers incudes flanking attB1 and attB2 Gateway recombination sites and the product was recombined first into pDONR201 and then into pPW-attB (gift of Mike Buszczak), an untagged UASp vector from the Drosophila Gateway collection modified to include an phiC-31 attB recombination target. The pUASp-Kelch^ΔNTR^ was injected into a strain carrying the attP2 phiC-31 landing site on Chromosome 3L at Rainbow transgenics.

#### Yeast two-hybrid screen

The Kelch KREP domain was cloned into pEG202, the bait vector for the LexA yeast two-hybrid system (Golemis et al., 2011). For the interaction screen, 6 × 106 yeast transformants from the Ovo II ovarian 2-Hybrid cDNA library (Grosshans et al., 1999) were screened for interaction by *β*-galactosidase production on X-gal plates and by complementation of leu2 auxotrophy. 200 of the fastest growing colonies were selected for further analysis.

#### CRISPR/Cas9 mutagenesis of *hts*

Oligos encoding sgRNAs described in Reagents Table were cloned into the BbsI site of the pBFvU6.2 vector (Kondo and Ueda, 2013) and integrated at the attP2 or attP40 phiC31 integration site (Groth et al., 2004). Mutagenesis was carried out in males expressing Cas9 from a germline promoter (nanos-Cas9; Port et al. (2014); Bloomington #54591) in combination with ubiquitous sgRNA expression from the snRNA:U6:96Ab promoter. Single male progeny bearing CRISPR-mutagenized chromosomes were crossed to a deficiency for *hts* (*Df(2R)BSC135/CyO*, Bloomington #9423), and hemizygous progeny were screened by sequencing and/or fertility tests. Balanced stocks were established for mutations of interest.

#### Construction of TagRFP-T::Ovhts::Venus BAC transgene

BAC clone CH321-84O22 (Venken et al., 2009), which contains all of the *hts* locus on a 91 kb genomic fragment (chr2R:19,368,835..19,460,115, FlyBase release 6), was used as a template to introduce fluorescent protein coding sequences. HA::Venus::FLAG sequence was inserted before the stop codon of exon 12, the exon encoding HtsRC (codon starting position chr2R:19,3399,860), using 2-step BAC recombineering. Briefly, a RpsL-neo selection cassette (Wang et al., 2009) was inserted using Kanamycin selection, and the RpsL-neo cassette was subsequently replaced with HA::Venus::FLAG using streptomycin counterselection. The same approach was then used to insert HA::TagRFP-T::FLAG before the first codon of *ovhts* exon 8 (Trp 473 of Ovhts polypeptide, codon starting position chr2R:19,409,361). Two additional rounds of recombineering were used to reduce the size of the BAC; the trimmed BAC contained a 44 kb genomic fragment from chr2R:19,390,835 (in CG11257) to chr2R:19,434,833 (in Fak). The BAC was injected into the attP2 landing site on chr3L at GenetiVision (Houston, TX). TagRFP-T fluorescence was not observed in transformed lines, suggesting that the introduction of TagRFP-T into the Adducin portion of Ovhts was detrimental to its folding. However, Venus fluorescence recapitulated the expression and localization patterns that are observed with the HtsRC monoclonal antibody, indicating that the cleaved ring canal protein tagged with Venus was produced normally. Consistent with this, the HtsRC protein produced by this construct was sufficient to rescue the loss of RC F-actin in *htsRC*-specific mutants, similar to otu-Ovhts::GFP in Figure S3.

#### Fixation, immunofluorescence, and imaging

Ovaries were fixed in 4% paraformaldehyde (Electron Microscopy Sciences) in PBS with or without 0.1% Triton X-100 for 10 minutes, washed in PBT (PBS, 0.1% Triton X-100, 0.5% BSA), and incubated in primary antibody in PBT at 4°C overnight. Antibodies and other fluorescent reagents are listed in Reagents Table. Samples were washed 4x in PBT and incubated with secondary antibodies and fluorescent phalloidin (if used) in PBT for 2 hours at room temperature. Following secondary antibody incubation, samples were washed 4x in PBT and mounted on slides in ProLong Gold or Diamond antifade reagents (Thermo Fisher Scientific). Samples were imaged on one of three microscopes: a Leica SP8 scanning confocal system using a 40x Plan Apo 1.30 NA objective, a Nikon TiE inverted microscope with a Yokogawa CSU-W1 spinning disc system, Andor iXon Ultra888 1024 × 1024 EMCCD, and 40x Plan Fluor 1.30 oil immersion objective, or a Zeiss Axiovert 200m inverted microscope with a CrEST X-light spinning disc system, Photometrics CoolSNAP HQ2 camera and either a 20x Plan Apo 0.8 NA objective or a 40x C-Apo 1.2 NA water-immersion objective. Images were processed and analyzed with ImageJ/FIJI or Imaris 9.0 (Bitplane).

#### Quantification of HtsRC immunolabeling at ring canals

Egg chambers were labeled with HtsRC 6A (Robinson et al., 1994) or Kel 1B monoclonal antibodies, both of which result in specific labeling of ring canals with little plasma membrane signal. Image stacks of wild-type and mutant egg chambers that included all ring canals in selected stage 10 egg chambers were collected on a spinning disc confocal with constant settings for exposure time, illumination power, and camera gain. Image stacks were analyzed in Imaris using automatic 3D thresholding of HtsRC or Kelch signal to identify ring canal volumes, and the summed fluorescence intensity was calculated for each ring canal. Raw fluorescent intensities were normalized to the mean fluorescence intensity of the control sample, which was defined as 1.

#### Quantification of ring canal F-actin thickness

To measure ring canal F-actin thickness, egg chambers were stained with fluorescent phalloidin. Maximum intensity projections of imaged egg chambers were rendered using Image J, and ring canals that provided a full longitudinal cross-section with a top-down view were analyzed (see top panel of Figure 3 for representative ring canals). A line was drawn through the center of measurable ring canals, and a F-actin fluorescence intensity plot spanning the line was generated. The full width at half maximum (FWHM) function of F-actin fluorescence intensity was calculated and used to approximate the thickness of the ring canal F-actin (see Hudson et al. (2015) for more details). Ring canals from stage 6 and stage 9 egg chambers were analyzed for Figure 3A and 3B, respectively. For Figure 3 graphs, n refers to the number of ring canals analyzed, the black line denotes the mean F-actin thickness, and error bars represent the 95% confidence interval. One-way ANOVA with Tukey’s multiple comparison correction was used in GraphPad Prism to detect for statistically significant differences in the mean ring canal F-actin thickness between samples.

#### Quantification of HtsRC fluorescent fusion protein levels

To quantify how HtsRC fluorescent fusion proteins levels were dependent on Kelch (Figure 4), egg chambers of all stages were imaged on a Zeiss spinning disc microscope using the same exposure time (500 ms). Maximum intensity projections of imaged egg chambers were rendered in Image J, and the maximum fluorescence value for each ring canal was calculated. The mean maximum fluorescence intensity of *Ovhts::Venus x2* sample (Figure 4B, black) was normalized to 100 relative fluorescence units. Each plotted point represents the maximum fluorescence intensity of the HtsRC fluorescent fusion protein (HtsRC::Venus for Figure 4B, HtsRC::GFP for Figure 4D). n equals the number of ring canals analyzed, and the black line represents the mean maximum fluorescence intensity while error bars represent 95% confidence interval. Statistical analyses (one-way ANOVA for Figure 4B and Student’s t-test for Figure 4D) were performed in GraphPad Prism to test for differences in the average maximum fluorescence intensity between the various genotypes, as indicated in the figure legend. Data are compiled from two independent experiments.

#### Western analysis of HtsRC protein levels from ovary extracts

To analyze HtsRC protein levels by western analysis, ovary lysates were generated by homogenizing dissected ovaries in SDS-PAGE sample buffer. One ovary equivalent was loaded per lane and separated on a 8.5 or 9% polyacrylamide gel. Proteins were transferred to nitrocellulose membrane, stained with amido black to visualize total protein, blocked in 5% milk in TBST (Tris-buffered saline with 0.1% Tween-20), and probed with the following antibodies: anti-HtsRC, anti-Kelch, anti-Tubulin. Blots were incubated with HRP-conjugated secondary antibodies followed by ECL development (Bio-Rad) and imaged on a CCD camera (Protein Simple). For Figure 5, levels of HtsRC, Kelch and *β*-tubulin were measured in FIJI/ImageJ, with *β*-tubulin serving as a loading control. Levels of HtsRC and Kelch are expressed relative to wild-type levels in *w*^*1118*^, defined as 1. Data are from four independent experiments. For Figure S4, total HtsRC protein levels were quantified using Image J by measuring the integrated density of HtsRC::GFP and HtsRC protein bands relative to amido black integrated density to control for protein levels loaded. HtsRC levels for mCherry-expressing control lysates were normalized to 1. HtsRC and HtsRC::GFP levels were parsed apart for graphical visualization. Error bars represent SEM. Student’s t-test was performed using GraphPad Prism. Data are from 3 independent experiments.

#### Analysis of HtsRC ubiquitylation and stability in S2 cells

S2 cells at 50-80% confluency were transfected with 1 *μ*g of pAHW-HtsRC::HA DNA following Effectene Transfection Reagent protocol (Qiagen). To assess HtsRC ubiquitylation and stability in response to drug/inhibitor treatments, cells were treated with DMSO control (Sigma-Aldrich), 1 *μ*M Bortezomib (Cell Signaling), and/or 100 *μ*g/mL Cycloheximide (Sigma-Aldrich) beginning at 24 or 36 hours post-transfection, as indicated in Figure 7 legend. Cells were harvested at the indicated time points (0, 3, 6 hours following inhibitor treatment), lysed in SDS-PAGE sample buffer, and analyzed by western analysis to visualize HtsRC and Tubulin protein levels. HtsRC protein levels were quantified using Image J by measuring the integrated density of HtsRC protein bands relative to Tubulin protein bands to control for protein loading. GraphPad Prism was used to perform a Student’s t-test using data from 3 independent experiments.

#### Quantification and Statistical Analysis

Data visualization and statistical analysis was performed in Prism (GraphPad). For quantification of protein levels by western blotting, error bars represent the mean *±* SEM. For all other quantification experiments, individual data points are plotted and the error bars represent the mean±95% confidence interval (CI), as indicated in the figure legend. Binomial test for enrichment of PEAEQ-containing proteins compared observed (15 PEAEQ proteins out of 141 KREP-interacting proteins) to expected (~3/141, based on the Drosophila genome encoding 262 PEAEQ proteins from 13931 protein coding genes (Table S1)). Other statistical tests used (one-way ANOVA or Student’s t-test), p values, and n are also listed in the figure legends.

## Supporting Information

### Reagents Table

#### List of Supplemental Tables

**Table S1**, related to Figure 1. List of proteins identified by mass spectrometry in KREP and control purifications.

#### List of Supplemental Figures

**Figure S1**, related to Figure 1. cDNA clones identified in yeast 2-hybrid screen with Kelch KREP domain contain PEAEQ-like sequences.

**Figure S2**, related to Figure 1. Conservation of *hts* PEAEQ motif among Drosophila species.

**Figure S3**, related to Figure 2. *ovhts* exon 12 truncation alleles specifically disrupt RCs and do not affect fusome.

**Figure S4**, related to Figure 4. HtsRC protein levels are dependent on Kelch by western analysis.

**Figure S5**, related to Figures 4 and 5. Expression levels of germline Gal4 drivers used in this study.

**Figure S6**, related to Figure 6. Analysis of HtsRC and Kelch levels in wild type, *hts*^Δ*Q925*^, and *hts*^Δ*E922*^.

#### REAGENTS TABLE

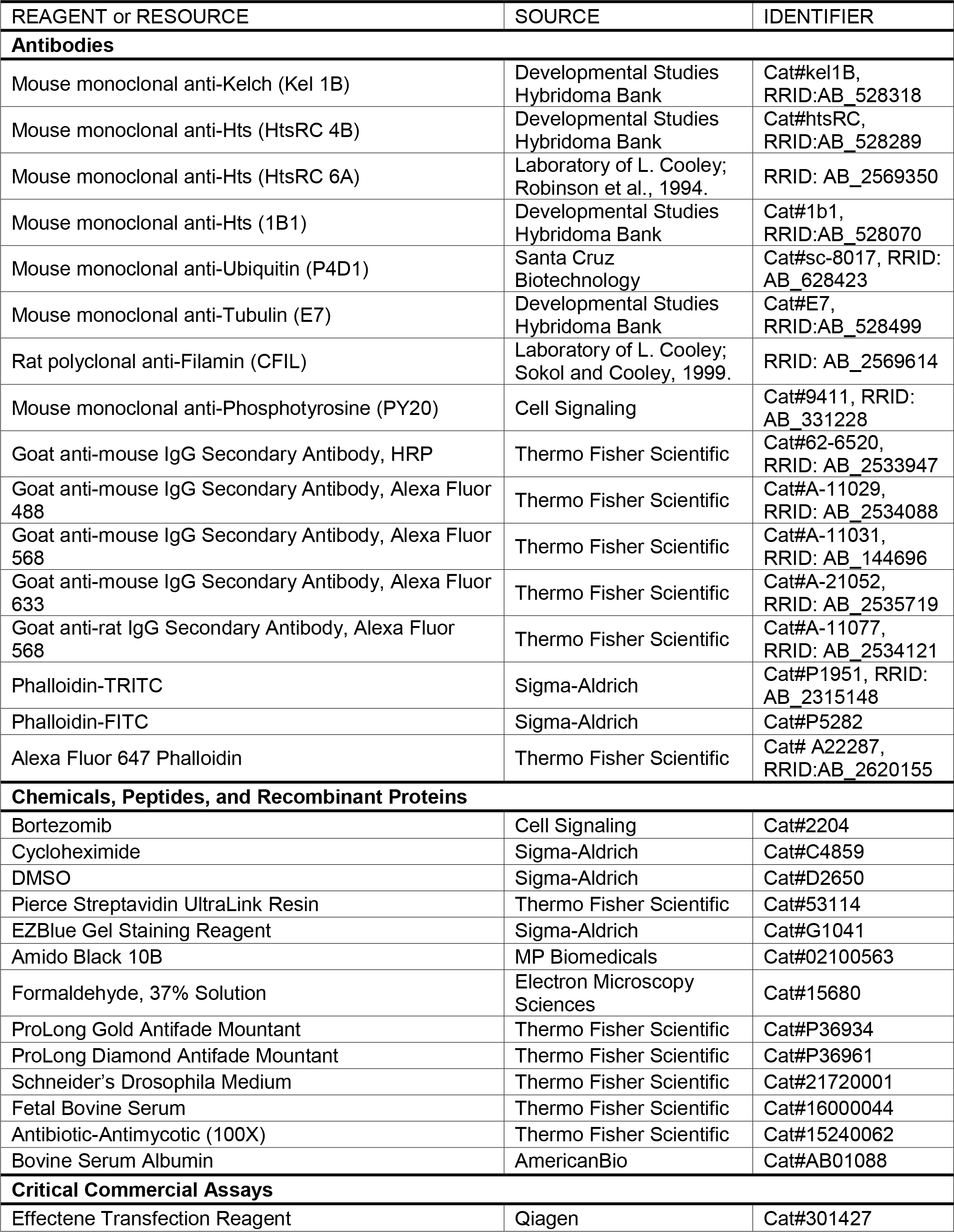

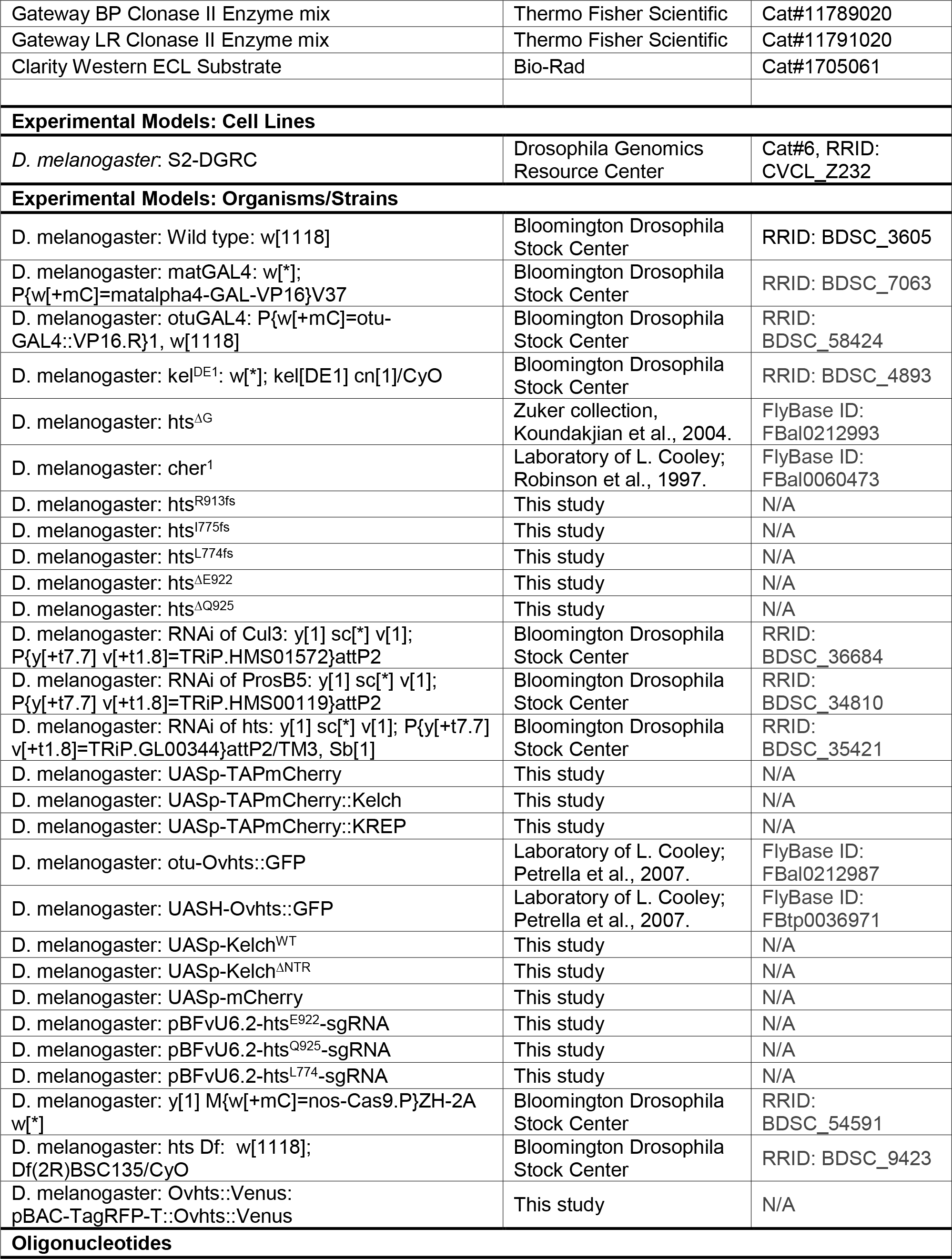

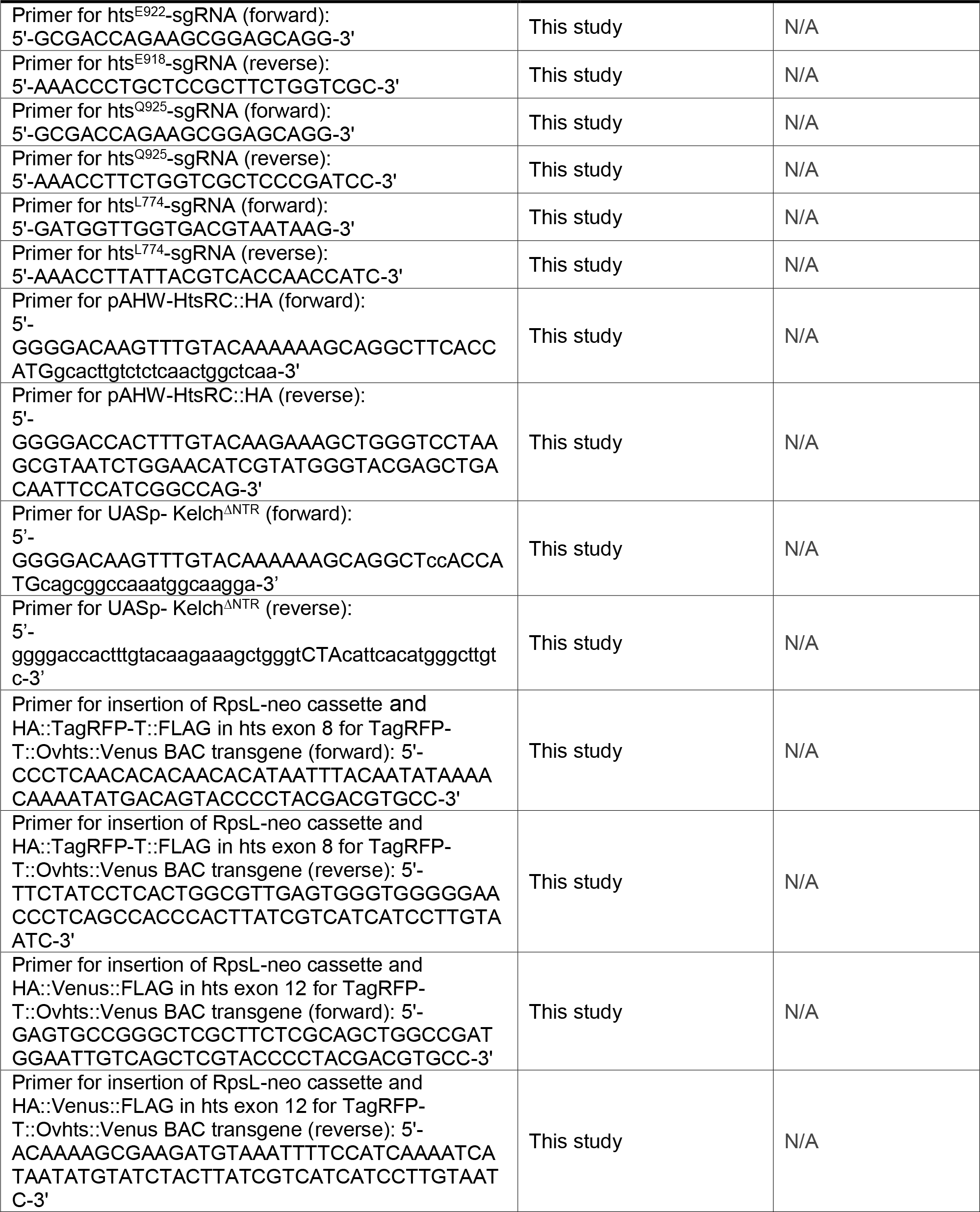

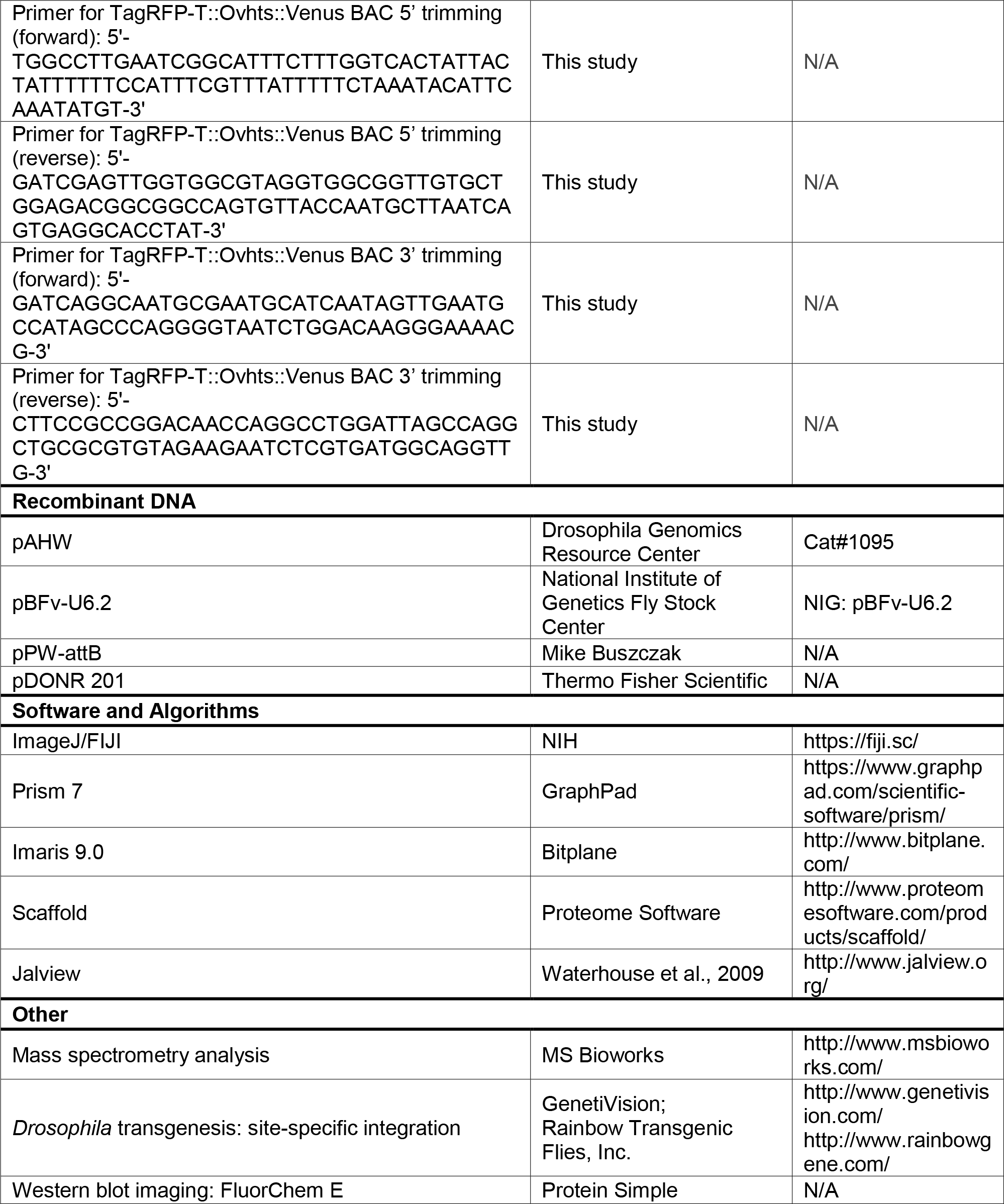

**Figure S1.**
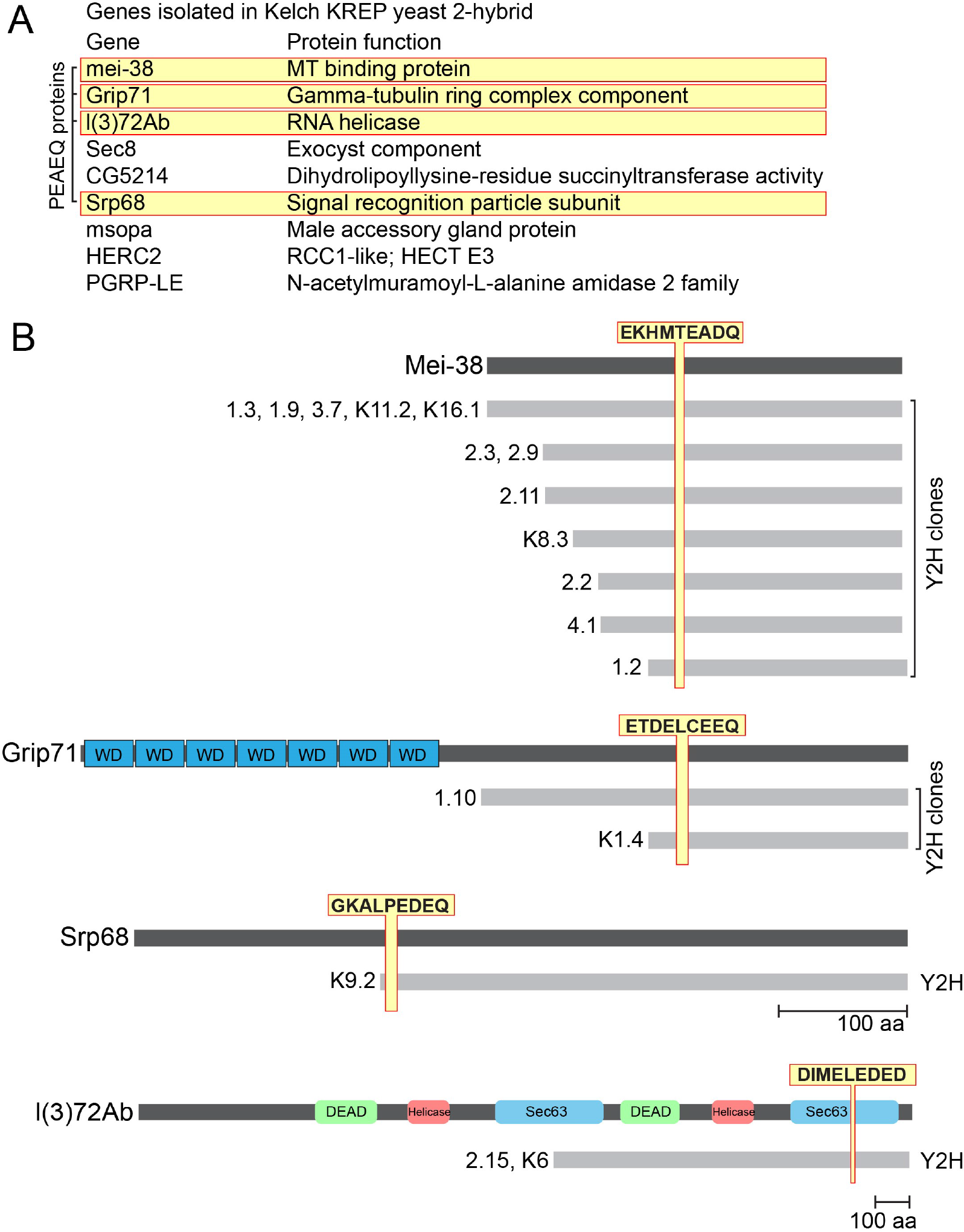
cDNA clones identified in yeast 2-hybrid screen with Kelch KREP domain contain PEAEQ-like sequences. (A) List of proteins identified in two hybrid screen with the KREP domain of Kelch as bait. Genes boxed in yellow contain PEAEQ motifs that satisfy the expression [PLT]-E-[AD]-[DE]-[QD]. (B) Diagrams of PEAEQ-containing proteins identified in screen; gray lines indicate extent of cDNA included in yeast 2-hybrid clones. In all cases, the cDNA clone included the sequence encoding the PEAEQ motif.

**Figure S2.**
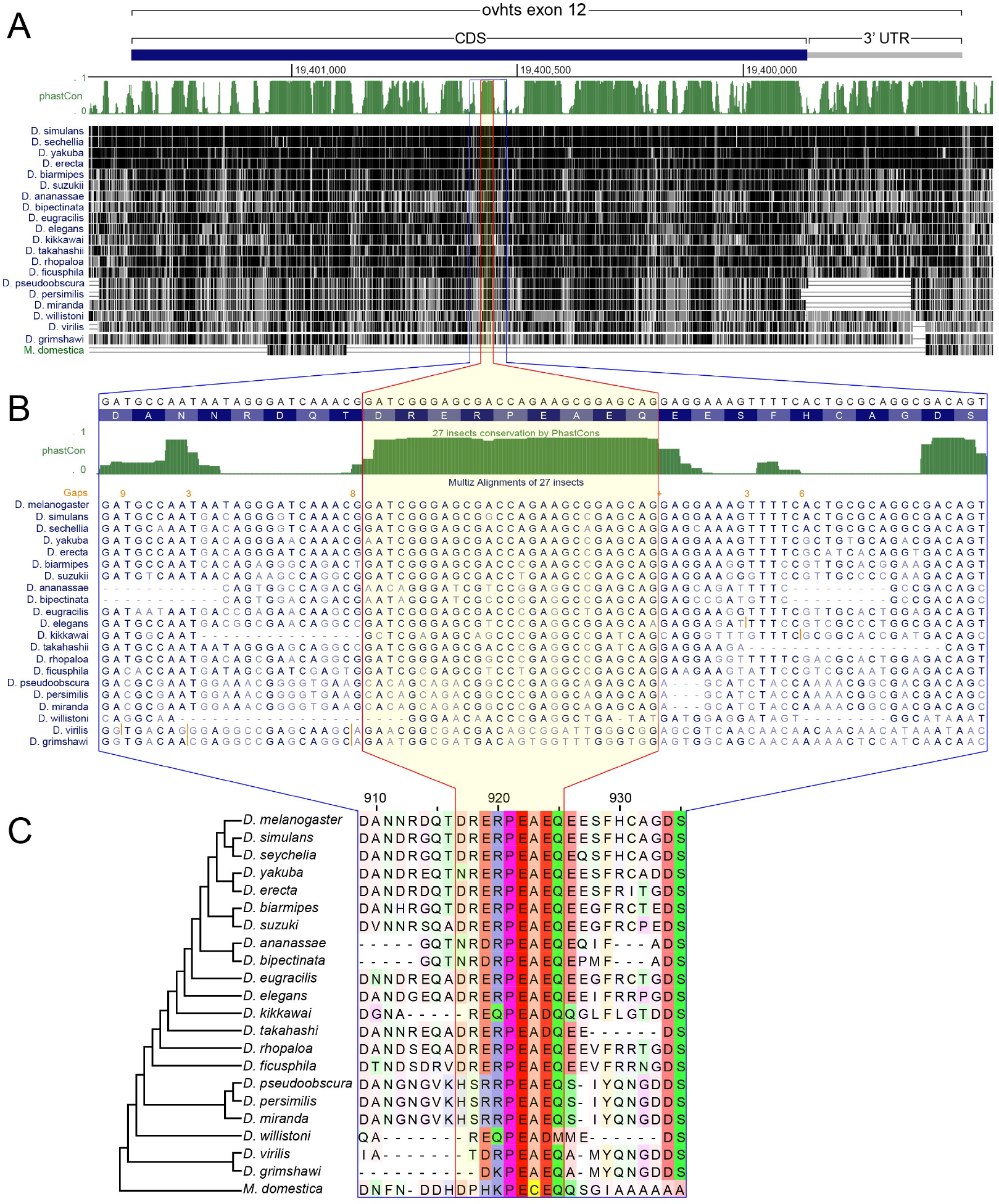
Conservation of *hts* PEAEQ motif among *Drosophila* species. (A) Exon 12 and the 3’ UTR of the *ovhts* transcript shown with graphical displays of nucleotide conservation derived from the PhastCons track of the UCSC genome browser. The PhastCons conservation track (green) shows the probability that given nucleotide is part of a conserved sequence element (Siepel et al., 2005). Individual sequence tracks of 20 *Drosophila* species are shown below with black shading indicating sequence conservation to the *melanogaster* reference genome. The sequence encoding the DRERPEAEQ element of HtsRC outlined in red is highly conserved in *Drosophila* species. The blue outline indicates the 81 nt that encode the DRERPEAEQ sequence in addition to less conserved flanking residues. (B) Nucleotide sequence conservation of 81 nt DRERPEAEQ region. Central region of high-scoring PhastCons sequence coincides with sequence encoding the DRERPEAEQ motif. (C) Alignment of peptide sequence encoded by the 81 nt region shown in B. Darker shading indicates higher conservation and reveals clear conservation of this motif in *Drosophila*. Alignments were constructed using the MAFFT program in Jalview (Waterhouse et al., 2009). In addition, the *Musca domestica* genome encodes an exon within its *hts* locus that corresponds to exon 12 of *Drosophila ovhts*. The *Musca* exon has diverged from the Drosophila sequence (bottom track in A), but a recognizable PEAEQ-like motif can be aligned to the Drosophila sequences

**Figure S3.**
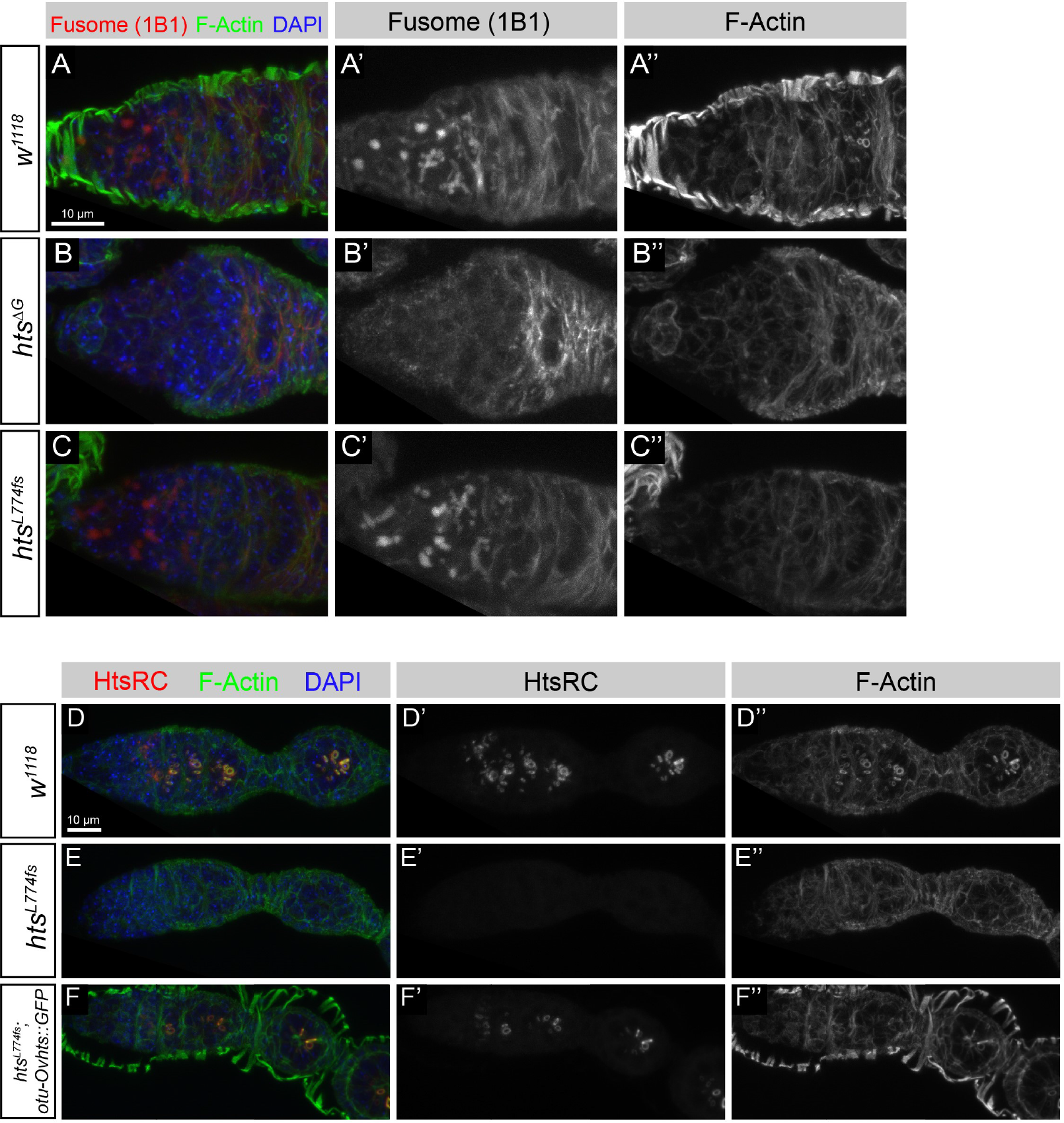
*ovhts* exon 12 truncation alleles specifically disrupt RCs and do not affect fusome. (A-C) Analysis of fusome integrity in wild type, *hts*^Δ*G*^, and *hts^L774fs^*. Wild-type egg germaria labeled with 1B1 antibody reveal HtsF protein on fusomes (A-A″). *hts*^Δ*G*^ is frameshift mutation that results in a nonsense codon just prior to the *ovhts* exon 12 splice site. It therefore cannot produce proteins with any of the C-terminal Adducin functional domains and behaves as a strong loss-of-function allele for all *hts* functions (Petrella et al., 2007). In *hts*^Δ*G*^, fusome labeling is undetectable in germaria labeled using 1B1 (B-B″). *hts*^*L774fs*^ mutations retain 1B1 HtsF labeling and do not exhibit the germline mitotic defects associated with fusome loss in *hts*^Δ*G*^ (C-C″). (D-F) HtsRC expression is eliminated in *hts*^*L774fs*^ mutants (E-E″), while expression of Ovhts::GFP restores HtsRC labeling and rescues the ring canal phenotype (F-F″).

**Figure S4.**
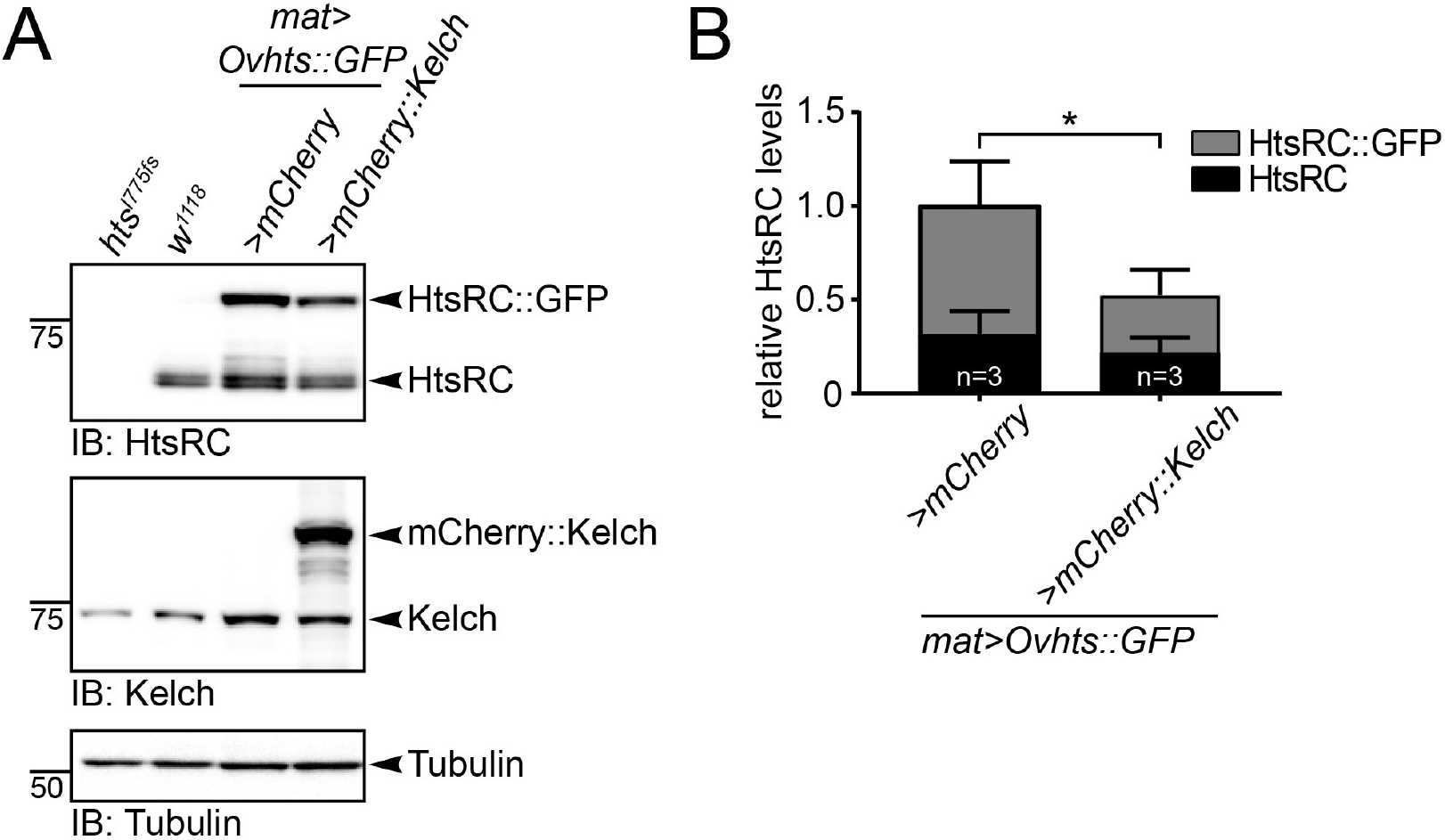
HtsRC protein levels are dependent on Kelch. (A) Representative western blot showing HtsRC, Kelch, and Tubulin protein levels from ovary lysates of indicated genotypes. (B) Quantification by western analysis of total HtsRC protein levels, normalized to Tubulin to control for protein loading. Levels of HtsRC::GFP and endogenous HtsRC species were parsed apart for graphical visualization. Data are from three independent experiments. Error bars indicate SEM. (*) p<0.05; Student’s t-test.

**Figure S5.**
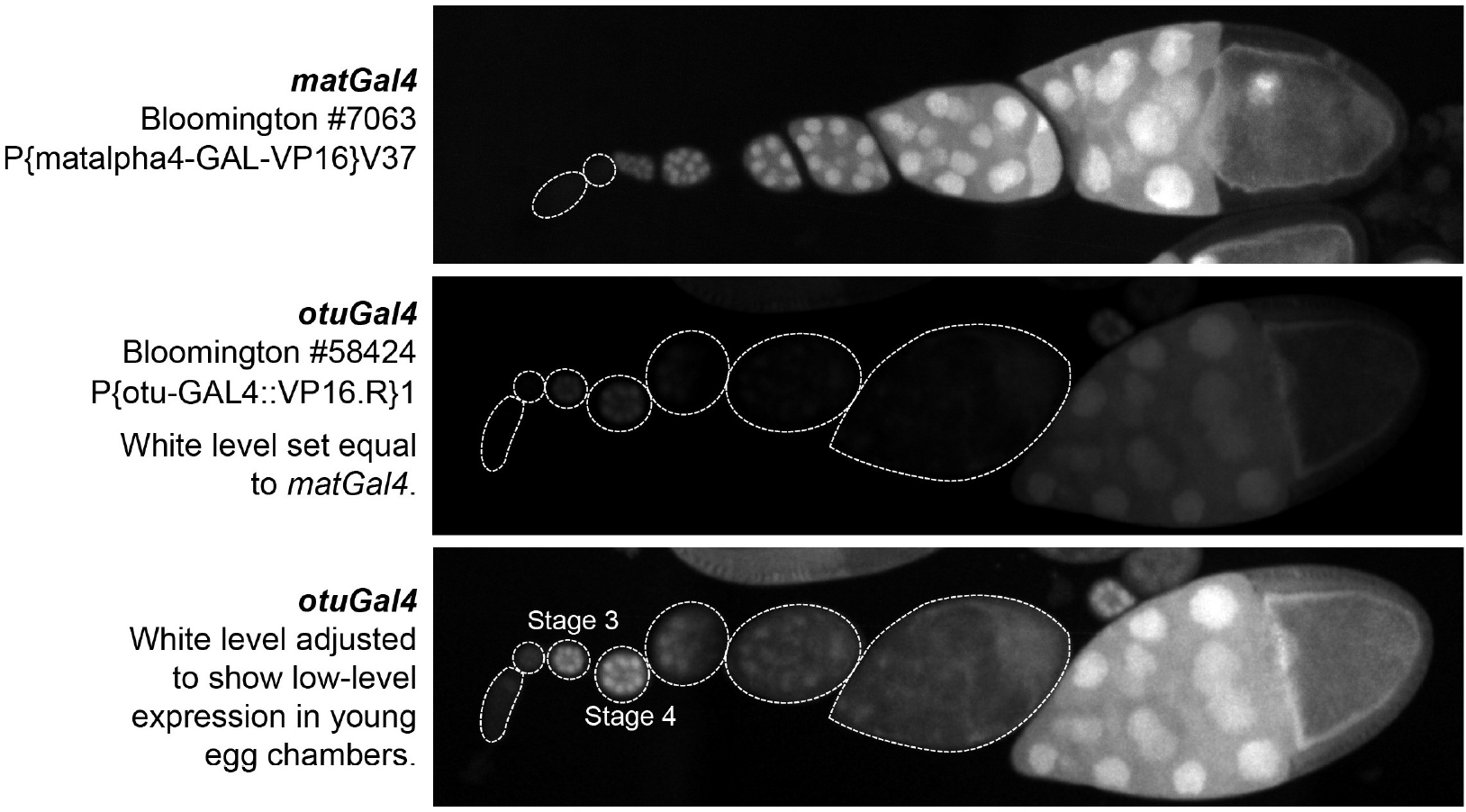
Expression levels of germline Gal4 drivers used in this study. Images of Gal4 driving expression of pUASp-mCherry are shown for *matGal4* and *otuGal4* drivers used in this study. *matGal4* drives little expression in the germarium, but then drives high-level expression in all subsequent stages. *otuGal4* drives expression at much lower levels, with detectable expression in stage 2-5 egg chambers, and then again in at stage 9-10.

**Figure S6.**
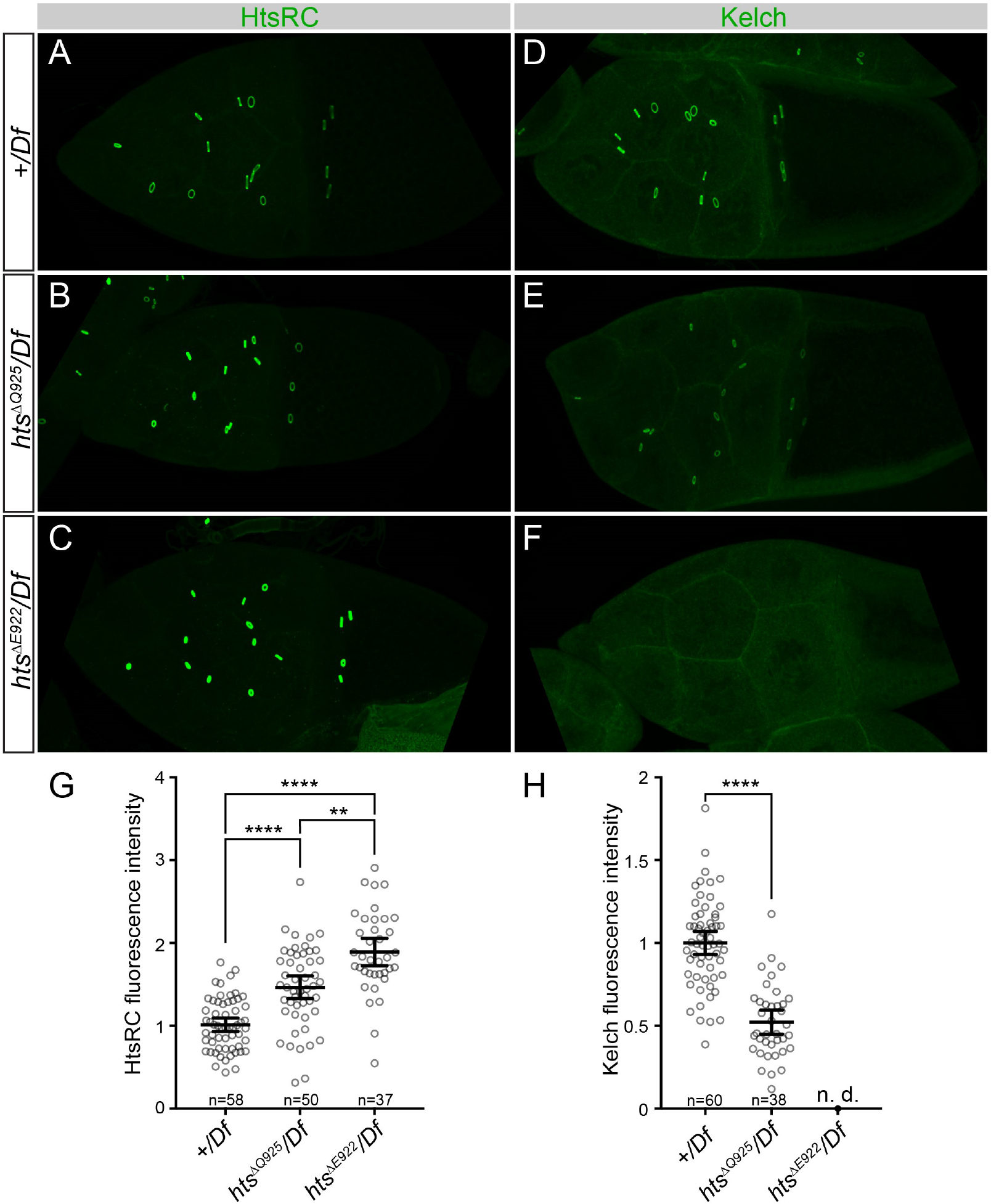
Analysis of HtsRC and Kelch levels in wild type, *hts*^Δ*Q925*^, and *hts*^Δ*E922*^. Single-codon deletion alleles, *hts*^Δ*Q925*^ and *hts*^Δ*E922*^, remove the conserved glutamine or glutamic acid codons in of the PEAEQ sequence. The *hts*^Δ*Q925*^ allele carried a linked lethal mutation, so this analysis compared the wild type, *hts*^Δ*Q925*^, and *hts*^Δ*E922*^ chromosomes in trans to *Df(2R)BSC256*, a deficiency that uncovers *hts* (abbreviated as *Df/+*). (A-C) Representative images of *Df*/+, *hts*^Δ*Q925*^/*Df* and *hts*^Δ*Q925*^/*Df* stage 10 egg chambers labeled for HtsRC. *hts*^Δ*Q925*^/*Df* mutants exhibit levels of HtsRC at ring canals that are intermediate between wild type and *hts*^Δ*E922*^/*Df*, suggesting that the ability of CRL3^Kelch^ to target HtsRC is reduced, but not eliminated. (G) Quantification of the HtsRC summed RC fluorescence intensities for the genotypes shown in A-C. The mean ring canal intensity of *Df*/+ was set at 1. Bars represent mean HtsRC fluorescence intensity ± 95% confidence interval. (****) p<0.0001; one-way ANOVA with Tukey’s multiple comparison test. (D-F) Representative images of *Df*/+, *hts*^Δ*Q925*^/*Df* and *hts*^Δ*E922*^/*Df* stage 10 egg chambers labeled for Kelch. Consistent with a reduction in CRL3^Kelch^ targeting efficiency in the *hts*^Δ*Q925*^/*Df* mutant, levels of Kelch are reduced, but not eliminated in *hts*^Δ*Q925*^/*Df* mutant compared to the *Df*/+ control. (H) Quantification of the Kelch summed ring canal fluorescence intensities for the genotypes shown in E-G. Bars represent mean Kelch fluorescence intensity ± 95% confidence interval. (****) p<0.0001; Student’s t-test. n.d., not determined.

## Acknowledgments

We thank Jovan Williams and Sowjanya Kallakuri for their help in recombineering the Ovhts BAC transgene. We thank Tian Xu for providing access to the Leica SP8 confocal microscope. Stocks obtained from the Bloomington Drosophila Stock Center (NIH P40OD018537) were used in this study. We thank the TRiP at Harvard Medical School for providing transgenic RNAi fly stocks used in this study. We thank the Drosophila Genomics Resource Center, supported by NIH grant 2P40OD010949, for the S2 cells used in this study. K.M.M. was supported in part by the National Institute of General Medical Sciences NIH training grant T32 GM007223. J.A.G. was funded in part by the National Institute of General Medical Sciences NIH training grant T32 GM007499 and a Gruber Science Fellowship. This work was funded by NIH grants: R01 GM043301 and RC1 GM091791.

## Author Contributions

Conceptualization, A.M.H., K.M.M., L.C.; Investigation, A.M.H., K.M.M., J.A.G., L.C.; Methodology, A.M.H., K.M.M., J.A.G., M.C.K., L.C.; Writing – Original Draft, A.M.H., K.M.M.; Writing – Review & Editing, A.M.H., K.M.M., J.A.G., M.C.K., L.C.; Visualization - A.M.H., K.M.M., J.A.G.; Funding Acquisition, L.C.; Resources, A.M.H., K.M.M., J.A.G., M.C.K., L.C.; Supervision, L.C

## Declaration of Interests

These authors have no competing interests.

## References

J. Adams, R. Kelso, and L. Cooley. The kelch repeat superfamily of proteins: propellers of cell function. Trends Cell Biol, 10:17–24, January 2000.

E. Arama, J. Agapite, and H. Steller. Caspase activity and a specific cytochrome C are required for sperm differentiation in Drosophila. Dev Cell, 4:687–97, May 2003.

E. Arama, M. Bader, G. E. Rieckhof, and H. Steller. A ubiquitin ligase complex regulates caspase activation during sperm differentiation in Drosophila. PLoS Biol, 5:e251, October 2007.

R. Bastock and D. St Johnston. Drosophila oogenesis. Curr Biol, 18:R1082–7, December 2008.

P. Bomont, L. Cavalier, F. Blondeau, C. Ben Hamida, S. Belal, M. Tazir, E. Demir, H. Topaloglu, R. Korinthenberg, B. Tuysuz, P. Landrieu, F. Hentati, and M. Koenig. The gene encoding gigaxonin, a new member of the cytoskeletal BTB/kelch repeat family, is mutated in giant axonal neuropathy. Nat Genet, 26:370–4, November 2000.

L. M. Boyden, M. Choi, K. A. Choate, C. J. Nelson-Williams, A. Farhi, H. R. Toka, I. R. Tikhonova, R. Bjornson, S. M. Mane, G. Colussi, M. Lebel, R. D. Gordon, B. A. Semmekrot, A. Poujol, M. J. Valimaki, M. E. De Ferrari, S. A. Sanjad, M. Gutkin, F. E. Karet, J. R. Tucci, J. R. Stockigt, K. M. Keppler-Noreuil, C. C. Porter, S. K. Anand, M. L. Whiteford, I. D. Davis, S. B. Dewar, A. Bettinelli, J. J. Fadrowski, C. W. Belsha, T. E. Hunley, R. D. Nelson, H. Trachtman, T. R. Cole, M. Pinsk, D. Bockenhauer, M. Shenoy, P. Vaidyanathan, J. W. Foreman, M. Rasoulpour, F. Thameem, H. Z. Al-Shahrouri, J. Radhakrishnan, A. G. Gharavi, B. Goilav, and R. P. Lifton. Mutations in kelch-like 3 and cullin 3 cause hypertension and electrolyte abnormalities. Nature, 482:98–102, February 2012.

M. R. Cummings, N. M. Brown, and R. C. King. The cytology of the vitellogenic stages of oogenesis in Drosophila melanogaster. 3. Formation of the vitelline membrane. Z Zellforsch Mikrosk Anat, 118:482–92, July 1971.

P. de Bie and A. Ciechanover. Ubiquitination of E3 ligases: self-regulation of the ubiquitin system via proteolytic and non-proteolytic mechanisms. Cell Death Differ, 18:1393–402, September 2011.

B. S. Dhanoa, T. Cogliati, A. G. Satish, E. A. Bruford, and J. S. Friedman. Update on the Kelch-like (KLHL) gene family. Hum Genomics, 7:13, May 2013.

E. A. Golemis, I. Serebriiskii, R. L. Finley, M. G. Kolonin, J. Gyuris, and R. Brent. Interaction trap/two-hybrid system to identify interacting proteins. Curr Protoc Cell Biol, Chapter 17: Unit 17 3, December 2011.

J. Grosshans, F. Schnorrer, and C. Nusslein-Volhard. Oligomerisation of Tube and Pelle leads to nuclear localisation of dorsal. Mech Dev, 81:127–38, March 1999.

A. C. Groth, M. Fish, R. Nusse, and M. P. Calos. Construction of transgenic Drosophila by using the site-specific integrase from phage phiC31. Genetics, 166:1775–82, April 2004.

V. A. Gupta and A. H. Beggs. Kelch proteins: emerging roles in skeletal muscle development and diseases. Skelet Muscle, 4:11, 2014.

K. Haglund, I. P. Nezis, and H. Stenmark. Structure and functions of stable intercellular bridges formed by incomplete cytokinesis during development. Commun Integr Biol, 4: 1–9, January 2011.

A. M. Hudson and L. Cooley. Drosophila Kelch functions with Cullin-3 to organize the ring canal actin cytoskeleton. J Cell Biol, 188:29–37, January 2010.

A. M. Hudson, K. M. Mannix, and L. Cooley. Actin Cytoskeletal Organization in Drosophila Germline Ring Canals Depends on Kelch Function in a Cullin-RING E3 Ligase. Genetics, 201:1117–31, November 2015.

Y. Kaplan, L. Gibbs-Bar, Y. Kalifa, Y. Feinstein-Rotkopf, and E. Arama. Gradients of a ubiquitin E3 ligase inhibitor and a caspase inhibitor determine differentiation or death in spermatids. Dev Cell, 19:160–73, July 2010.

D. Komander and M. Rape. The ubiquitin code. Annu Rev Biochem, 81:203–29, 2012.

Shu Kondo and Ryu Ueda. Highly Improved Gene Targeting by Germline-Specific Cas9 Expression in <em>Drosophila</em>. Genetics, 195:715–721, 2013.

E. J. Koundakjian, D. M. Cowan, R. W. Hardy, and A. H. Becker. The Zuker collection: a resource for the analysis of autosomal gene function in Drosophila melanogaster. Genetics, 167:203–6, May 2004.

W. Li, M. H. Bengtson, A. Ulbrich, A. Matsuda, V. A. Reddy, A. Orth, S. K. Chanda, S. Batalov, and C. A. Joazeiro. Genome-wide and functional annotation of human E3 ubiquitin ligases identifies MULAN, a mitochondrial E3 that regulates the organelle’s dynamics and signaling. PLoS One, 3:e1487, January 2008.

X. Li, D. Zhang, M. Hannink, and L. J. Beamer. Crystal structure of the Kelch domain of human Keap1. J Biol Chem, 279:54750–8, December 2004.

Z. Lin, S. Li, C. Feng, S. Yang, H. Wang, D. Ma, J. Zhang, M. Gou, D. Bu, T. Zhang, X. Kong, X. Wang, O. Sarig, Y. Ren, L. Dai, H. Liu, J. Zhang, F. Li, Y. Hu, G. Padalon-Brauch, D. Vodo, F. Zhou, T. Chen, H. Deng, E. Sprecher, Y. Yang, and X. Tan. Stabilizing mutations of KLHL24 ubiquitin ligase cause loss of keratin 14 and human skin fragility. Nat Genet, 48:1508–1516, December 2016.

S. C. Lo, X. Li, M. T. Henzl, L. J. Beamer, and M. Hannink. Structure of the Keap1:Nrf2 interface provides mechanistic insight into Nrf2 signaling. EMBO J, 25:3605–17, August 2006.

N. Matova and L. Cooley. Comparative aspects of animal oogenesis. Dev Biol, 231:291–320, March 2001.

E. Morais-de Sa, A. Vega-Rioja, V. Trovisco, and D. St Johnston. Oskar is targeted for degradation by the sequential action of Par-1, GSK-3, and the SCF(-)Slimb ubiquitin ligase. Dev Cell, 26:303–14, August 2013.

B. Padmanabhan, K. I. Tong, T. Ohta, Y. Nakamura, M. Scharlock, M. Ohtsuji, M. I. Kang, A. Kobayashi, S. Yokoyama, and M. Yamamoto. Structural basis for defects of Keap1 activity provoked by its point mutations in lung cancer. Mol Cell, 21:689–700, March 2006.

J. Pae, R. M. Cinalli, A. Marzio, M. Pagano, and R. Lehmann. GCL and CUL3 Control the Switch between Cell Lineages by Mediating Localized Degradation of an RTK. Dev Cell, 42:130–142 e7, July 2017.

J. S. Peterson, A. K. Timmons, A. A. Mondragon, and K. McCall. The End of the Beginning: Cell Death in the Germline. Curr Top Dev Biol, 114:93–119, 2015.

L. N. Petrella, T. Smith-Leiker, and L. Cooley. The Ovhts polyprotein is cleaved to produce fusome and ring canal proteins required for Drosophila oogenesis. Development, 134: 703–12, February 2007.

F. Port, H. M. Chen, T. Lee, and S. L. Bullock. Optimized CRISPR/Cas tools for efficient germline and somatic genome engineering in Drosophila. Proc Natl Acad Sci U S A, 111: E2967–76, July 2014.

S. Prag and J. C. Adams. Molecular phylogeny of the kelch-repeat superfamily reveals an expansion of BTB/kelch proteins in animals. BMC bioinformatics, 4:42, 2003.

D. N. Robinson and L. Cooley. Drosophila kelch is an oligomeric ring canal actin organizer. J Cell Biol, 138:799–810, August 1997.

D. N. Robinson, K. Cant, and L. Cooley. Morphogenesis of Drosophila ovarian ring canals. Development, 120:2015–25, July 1994.

D. N. Robinson, T. A. Smith-Leiker, N. S. Sokol, A. M. Hudson, and L. Cooley. Formation of the Drosophila ovarian ring canal inner rim depends on cheerio. Genetics, 145:1063–72, April 1997.

E. C. Roosen-Runge. Comparative aspects of spermatogenesis. Biol Reprod, 1:Suppl 1:24–31, June 1969.

F. R. Schumacher, F. J. Sorrell, D. R. Alessi, A. N. Bullock, and T. Kurz. Structural and biochemical characterization of the KLHL3-WNK kinase interaction important in blood pressure regulation. Biochem J, 460:237–46, June 2014.

S. Shibata, J. Zhang, J. Puthumana, K. L. Stone, and R. P. Lifton. Kelch-like 3 and Cullin 3 regulate electrolyte homeostasis via ubiquitination and degradation of WNK4. Proc Natl Acad Sci U S A, 110:7838–43, May 2013.

S. Shibata, J. P. Arroyo, M. Castaneda-Bueno, J. Puthumana, J. Zhang, S. Uchida, K. L. Stone, T.T. Lam, and R. P. Lifton. Angiotensin II signaling via protein kinase C phosphorylates Kelch-like 3, preventing WNK4 degradation. Proc Natl Acad Sci U S A, 111:15556–61, October 2014.

A. Siepel, G. Bejerano, J. S. Pedersen, A. S. Hinrichs, M. Hou, K. Rosenbloom, H. Clawson, J. Spieth, L. W. Hillier, S. Richards, G. M. Weinstock, R. K. Wilson, R. A. Gibbs, W. J. Kent, W. Miller, and D. Haussler. Evolutionarily conserved elements in vertebrate, insect, worm, and yeast genomes. Genome Res, 15:1034–50, August 2005.

N. S. Sokol and L. Cooley. Drosophila filamin encoded by the cheerio locus is a component of ovarian ring canals. Curr Biol, 9:1221–30, November 1999.

K. J. Venken, J. W. Carlson, K. L. Schulze, H. Pan, Y. He, R. Spokony, K. H. Wan, M. Koriabine, P. J. de Jong, K. P. White, H. J. Bellen, and R. A. Hoskins. Versatile P[acman] BAC libraries for transgenesis studies in Drosophila melanogaster. Nat Methods, 6:431–4, June 2009.

S. Wang, Y. Zhao, M. Leiby, and J. Zhu. A new positive/negative selection scheme for precise BAC recombineering. Mol Biotechnol, 42:110–6, May 2009.

A. M. Waterhouse, J. B. Procter, D. M. Martin, M. Clamp, and G. J. Barton. Jalview Version 2–a multiple sequence alignment editor and analysis workbench. Bioinformatics, 25: 1189–91, May 2009.

L. Xu, Y. Wei, J. Reboul, P. Vaglio, T. H. Shin, M. Vidal, S. J. Elledge, and J. W. Harper. BTB proteins are substrate-specific adaptors in an SCF-like modular ubiquitin ligase containing CUL-3. Nature, 425:316–21, September 2003.

F. Xue and L. Cooley. kelch encodes a component of intercellular bridges in Drosophila egg chambers. Cell, 72:681–93, March 1993.

D. Yan, R. A. Neumuller, M. Buckner, K. Ayers, H. Li, Y. Hu, D. Yang-Zhou, L. Pan, X. Wang, C. Kelley, A. Vinayagam, R. Binari, S. Randklev, L. A. Perkins, T. Xie, L. Cooley, and N. Perrimon. A regulatory network of Drosophila germline stem cell self-renewal. Dev Cell, 28:459–73, February 2014.

A. N. Yatsenko, A. Roy, R. Chen, L. Ma, L. J. Murthy, W. Yan, D. J. Lamb, and M. M. Matzuk. Non-invasive genetic diagnosis of male infertility using spermatozoal RNA: KLHL10 mutations in oligozoospermic patients impair homodimerization. Hum Mol Genet, 15: 3411–9, December 2006.

L. Yue and A. C. Spradling. hu-li tai shao, a gene required for ring canal formation during Drosophila oogenesis, encodes a homolog of adducin. Genes Dev, 6:2443–54, December 1992.

